# DEK oncoprotein participates in heterochromatin replication via SUMO-dependent nuclear bodies

**DOI:** 10.1101/2023.03.09.529154

**Authors:** Agnieszka Pierzynska-Mach, Christina Czada, Christopher Vogel, Eva Gwosch, Xenia Osswald, Denis Bartoschek, Alberto Diaspro, Ferdinand Kappes, Elisa Ferrando-May

## Abstract

The correct inheritance of chromatin structure is key for maintaining genome function and preventing cellular transformation. DEK, a conserved chromatin protein, has recognized tumor-promoting properties, its overexpression being associated with poor prognosis in various cancer types. At the cellular level, DEK displays pleiotropic functions, influencing differentiation, apoptosis, and stemness, but a characteristic oncogenic mechanism remains elusive. Here, we report the identification of DEK bodies, focal assemblies of DEK occurring at specific, yet unidentified sites of heterochromatin replication. In these bodies, DEK localizes in direct proximity to active replisomes suggesting a function in the early maturation of heterochromatin. A high-throughput siRNA screen identifies SUMO as a major regulator of DEK body formation, linking DEK to the SUMO network that controls chromatin states and cell fate. This work combines and refines our previous data on DEK as a factor essential for heterochromatin integrity and facilitating replication under stress and delineates an avenue of further study for unraveling DEK’s contribution to cancer development.

## Introduction

Replication of genomes is a fundamental process requiring meticulously orchestrated, yet adaptable mechanisms in space and time. The replication machinery encounters a range of different chromatin structures, some of which are prone to delaying, impairing, or even stalling replication fork progression - a state referred to as DNA replication stress (Maya Mendoza et al., 2018) and often encountered in cancer cells (Boyer et al., 2016; Saxena and Zou, 2022). Overcoming such obstacles requires the coordinated action of a number of protective chromatin factors, often governed by targeted post-translational modifications, with poly(ADP)ribosylation and SUMOylation playing prominent roles (Abbas, 2021; Stewart-Morgan et al., 2020). Active DNA replication forks, as visualized by incorporation of DNA precursors (BrdU, EdU or other thymidine analogues) or *via* marking DNA replication factors (e.g. Proliferating Cell Nuclear Antigen, PCNA) (Chagin et al., 2016; Jackson and Pombo, 1998) appear as a characteristic pattern of nuclear foci that changes over time and typically reflects the chromatin structure being replicated (Aladjem, 2007; Casas-Delucchi et al., 2012; Heinz et al., 2018; O’Keefe et al., 1992; Sehwaiger et al., 2009). In general, euchromatin undergoes DNA replication during early S-phase, facultative heterochromatin during middle S-phase, and constitutive heterochromatin during late S-phase (Chadwick and Willard, 2004; Fragkos et al., 2015; Ryba et al., 2010; Su et al., 2020). However, some “hard-to-replicate” DNA structures, e.g., inactivated X chromosome, telomeres, centromeres, or certain DNA secondary structures, such as G-quadruplexes, four-way junction DNA and others, pose a particular challenge to the DNA replication machinery, and, in turn, require fine-tuned action of additional accessory factors (Mirkin and Mirkin, 2007).

We and others have identified the DEK oncogene, a unique and multifunctional non-histone chromosomal protein, as a supportive factor of DNA replication, particularly in scenarios where the replication machinery is under stress (Deutzmann et al., 2015; Ganz et al., 2019). Although Isolation of Proteins on Nascent DNA (iPOND) studies support no direct association of DEK with the replisome, DEK is enriched in maturing chromatin (Alabert et al., 2014; Aranda et al., 2014; Garcia et al., 2017; Lossaint et al., 2013; Ribeyre et al., 2016; Sirbu et al., 2013), in line with its post-translational modification-dependent function as histone chaperone and its role in the maintenance of chromatin structure (Alexiadis et al., 2000; Böhm et al., 2005; Cavellán et al., 2006; Gamble and Fisher, 2007; Guo et al., 2021a; Hollenbach et al., 2002; Kappes et al., 2001; Kappes et al., 2004a; Kappes et al., 2004b; Kappes et al., 2011a; Privette Vinnedge et al., 2013; Saha et al., 2013; Sawatsubashi et al., 2010; Tabbert et al., 2006; Waidmann et al., 2014; Waldmann et al., 2002; Waldmann et al., 2003). Under moderate to severe replication stress, DEK is highly enriched at maturing chromatin, suggesting a protective role at the replication fork. Our previous research demonstrated that down-regulation of DEK in cells under moderate short-term replication stress *via* aphidicolin and camptothecin treatment results in the aggravation of replication stress phenotypes. Replication fork progression, proliferative capability and clonogenic survival were impaired and cells accumulated replication-born DNA damage and transmitted it to daughter cells (Deutzmann et al., 2015). Such fork-protective properties of DEK are modulated by PARP1/2 activity depending on the type and extent of replication stress (Ganz et al., 2019). At the same time, DEK appears to safeguard DNA replication also under steady-state, unchallenged conditions, as DNA fiber assays showed a mild decrease in fork speed when DEK expression is downregulated in untreated cells (Deutzmann et al., 2015). Collectively, these findings suggest that DEK is not a direct component of the replisome but binds in the vicinity of the fork to facilitate fork progression by supporting chromatin maturation. Evidence for a role of DEK in DNA replication via its properties as a chromatin architectural protein comes also from *in vitro* studies: experiments with SV40 mini chromosomes showed that DEK introduces positive supercoils to chromatin and reduces the replication efficiency of chromatin in a reconstituted system (Alexiadis et al., 2000; Waldmann et al., 2002). Furthermore, DEK preferentially associates with four-way junctions over duplex DNA and with supercoiled over linear DNA, both structures that can induce replication stress *per se* or can be consequences of replication stress (Böhm et al., 2005; Waldmann et al., 2003).

Under steady-state conditions DEK associates with chromatin throughout the cell cycle (Kappes et al., 2001; Matrka et al., 2015), typically observed as pan-nuclear punctate pattern in immunofluorescence images and further supported by the requirement for medium to high salt concentrations for its extraction from nuclei preparations (Kappes et al., 2001; Kappes et al., 2004a; Kappes et al., 2008; Özçelik et al., 2022; Saha et al., 2013), all in line with its pleiotropic involvement in multiple nuclear processes, ranging from transcriptional regulation, DNA damage response and repair, apoptosis, cell stemness and chromatin maintenance (Privette Vinnedge et al., 2013; Waldmann et al., 2004). More detailed analyses revealed that DEK can associate both with euchromatin and heterochromatin (Capitano et al., 2022; Gamble and Fisher, 2007; Guo et al., 2021a; Hollenbach et al., 2002; Hu et al., 2005; Ivanauskiene et al., 2014; Kappes et al., 2001; Kappes et al., 2004a; Kappes et al., 2004b; Kappes et al., 2008; Kappes et al., 2011b; Karam et al., 2014; Privette Vinnedge et al., 2013; Saha et al., 2013; Sandén and Gullberg, 2015; Sandén et al., 2014; Sawatsubashi et al., 2010; Waidmann et al., 2014; Zhou et al., 2022). However, qualitative and quantitative differences in the specific chromatin distribution of DEK amongst cell types and conditions are evident. Whereas DEK was identified as a suppressor of variegation (Su(var)), thus as a factor important for maintenance of heterochromatin (Capitano et al., 2019; Kappes et al., 2011a; Saha et al., 2013), other studies show preferential association with euchromatin (Hu et al., 2007; Ivanauskiene et al., 2014; Sandén et al., 2014; Yang et al., 2022), pinpointing to cell-type-specific post-translational modifications on DEK as critical for its specific chromatin association. Indeed, DEK is targeted by a plethora of post-translational modifications, with biological effects for acetylation, phosphorylation and poly(ADP)ribosylation having been reported (Cleary et al., 2005; Fahrer et al., 2010; Kappes et al., 2004a; Kappes et al., 2004b; Kappes et al., 2008; Mor-Vaknin et al., 2011; Tabbert et al., 2006). Moreover, DEK contains an RNA binding motif and is found associated with cellular RNA (Baltz et al., 2012; Beckmann et al., 2015; Castello et al., 2012; Guo et al., 2021a; Guo et al., 2021b). These data delineate DEK as a multifaceted protein subject to complex regulation and impacting multiple nuclear processes.

To deepen our insight in DEK′s functional mechanisms, in particular related to DNA replication, we investigated the dynamic localization of DEK in the nucleus throughout the cell cycle by live and high-resolution microscopy including optical nanoscopy (Diaspro and Bianchini, 2020). We discovered that, in late S-phase, DEK consistently forms discrete foci, now termed DEK bodies, which locate in close proximity of active heterochromatic replication sites. We then conducted an imaging-based siRNA screen to identify cellular regulators involved in the formation of DEK bodies. Besides methyltransferases, acetylases and proteins involved in checkpoint regulation, we surprisingly identified SUMO and ubiquitin enzymes, with SUMO1/3 as top candidates. Driven by this finding, we show that DEK is targeted by SUMOylation *in vitro* and in cells, and that SUMO modification of DEK is required, at least in part, for the establishment of DEK bodies. Our data refine our previous knowledge about DEK′s involvement in DNA replication and put forward DEK as a co-factor in the duplication of specific heterochromatic regions including pericentric chromatin. These findings provide further insights into the post-translational-dependent mechanisms by which DEK contributes to heterochromatin formation and maintenance with consequences for a variety of cellular processes.

## Results

### A sub-population of DEK molecules assembles into complex nuclear bodies at sites of late DNA replication

To obtain precise insights into the distribution of DEK on chromatin in the context of DNA replication, we first investigated its sub-nuclear localization throughout the cell cycle along with EdU or PCNA as replication fork markers. Confocal imaging of endogenous DEK in EdU-labeled MCF10A cells (**Fig. 1A**) revealed an archetypical, mostly uniform and punctuate nuclear pattern of DEK in G1/G2 and early to mid S-phase, in full agreement with previous observations as outlined above. In late S-phase, however, as far undescribed, distinct focal assemblies of DEK became visible which located in spatial proximity or within DNA replication foci as assessed either by EdU pulse labeling or by PCNA immunofluorescence (**Fig.1B, C**). We made the same observation in wild-type U2- OS cells **(Fig. 1D upper row**) and primary BJ-5ta cells (**Fig. S1A, B, C**), where S-phase was assessed by PCNA staining. Interestingly, in contrast to non-malignant MCF10A cells, no or very few foci occurred under identical conditions in MCF7 (non-invasive and poorly aggressive with low metastatic potential) or MDA-MB-231 (invasive metastatic) breast cancer cells (**Fig. S1D, E**), suggesting cell-type dependent differences and a potential correlation of DEK foci formation with malignancy. Focal accumulations of DEK in late S-phase were observed also with a previously established U2-OS cell line carrying a TALEN-mediated knock-in of eGFP-DEK, U2-OS KI eGFP-DEK cells (Ganz et al., 2019) (**Fig. 1D lower panel, Fig. S2A, B)** and when DEK was expressed transiently in U2-OS cells (**Fig. S2C**). Such accumulations were not detected in eGFP-only expressing cells **(Fig. S2D)**. Thus, formation of late replication-associated foci is a specific feature of DEK, possibly regulated by cellular state, for which the U2-OS KI eGFP-DEK cell line is a valid reporter. We term these foci henceforth “DEK bodies”.

**Figure 1:**
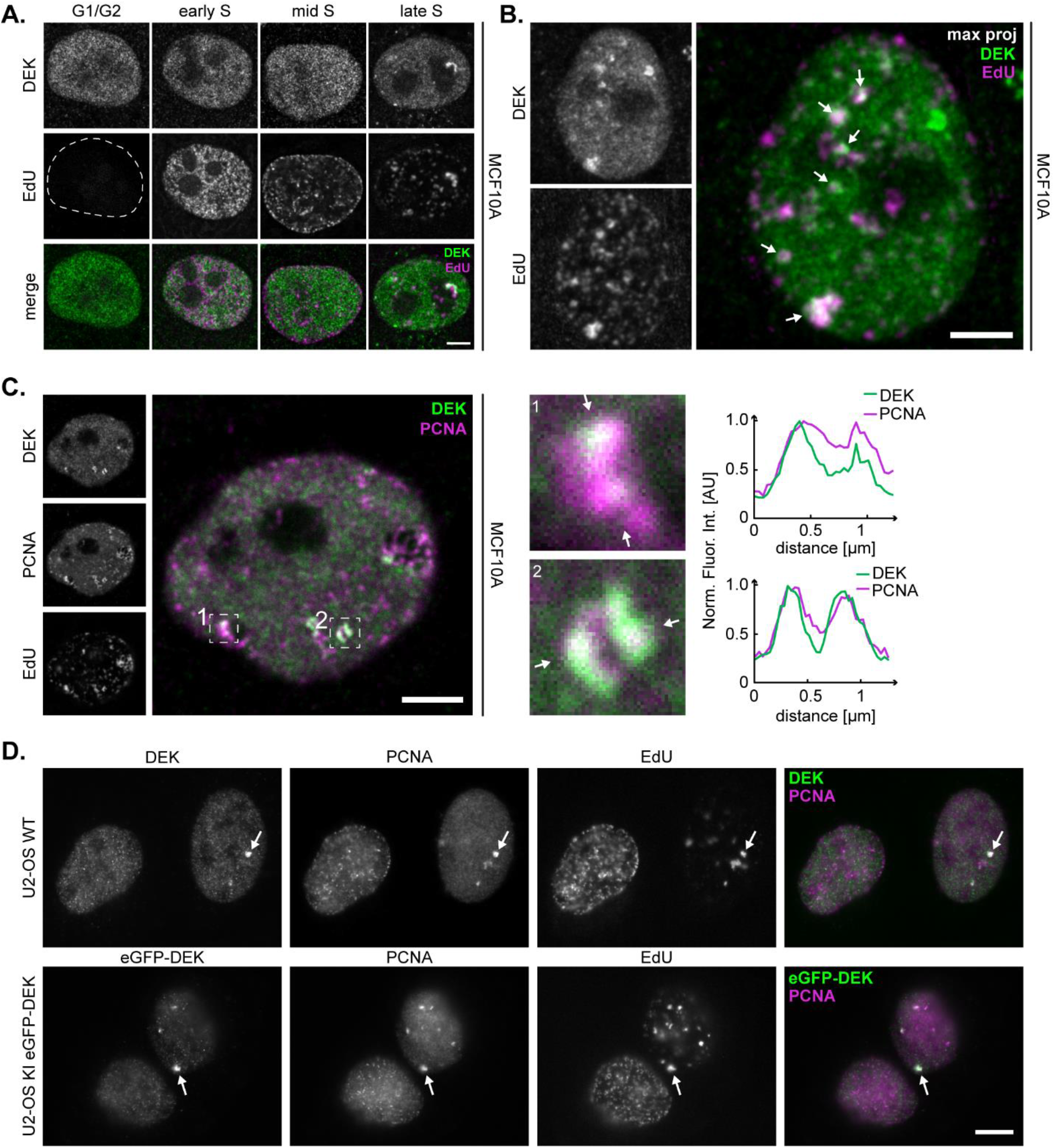
DEK forms distinct foci, DEK bodies, which colocalize with nascent DNA markers. **A.** Widefield images of MCF10A cells. Nascent DNA was pulse-labeled with EdU for 25 min followed by detection via click chemistry and immunofluorescence staining using DEK-specific antibodies. Average number of DEK foci per cell: 6.7 ± 2.2 (*n* = 46 cells). Magenta: EdU. Green: DEK. Scale bar: 5 μm. **B.** Immunofluorescence analysis of DEK and EdU distribution pattern in late S-phase MCF10A cells as described in (A) (maximum projection). Arrows point to sites of DEK and EdU colocalization. Scale bar: 5 μm. **C.** Immunofluorescence and colocalization analysis of DEK (green) and PCNA (magenta) in a late S-phase MCF10A cell nucleus. Dashed boxes indicate two selected ROIs. The intensity profiles are measured along the lines shown in the enlarged ROIs. The fluorescence intensity was normalized to 1. Scale bars: 5 μm. **D.** Widefield fluorescence images of immunolabelled U2-OS wild type cells (upper row) and U2-OS KI eGFP-DEK cells (bottom row). Endogenous DEK (green) and PCNA (magenta) were visualized by specific antibodies. In each image one prominent DEK body is marked by a white arrow. Scale bar: 5 μm.

To get deeper structural insight into DEK bodies and their spatial relation to active replication sites we turned to super-resolution imaging. We visualized firstly the granular fine structure of endogenous DEK bodies in MCF10A cells by Stimulated Emission Depletion (STED) and single-molecule localization (SML) microscopy (**Fig. 2A, B**). Secondly, we investigated the organization of nascent DNA and replication forks within DEK bodies of wild-type U2-OS cells by labeling with EdU and PCNA-specific antibodies followed by 3D Structured Illumination Microscopy (3D-SIM). The data show that DEK, PCNA, and EdU signals do not colocalize but are juxtaposed within DEK bodies (**Fig. 2C, D**). Intensity line plots (**Fig. 2E**) highlight the close spatial proximity, which was also observed in primary BJ-5ta foreskin fibroblasts (**Fig. S1**). This result is in line with previous data showing that DEK is not part of the replisome but binds to nascent chromatin in the vicinity of the replication fork (Ganz et al., 2019).

**Figure 2:**
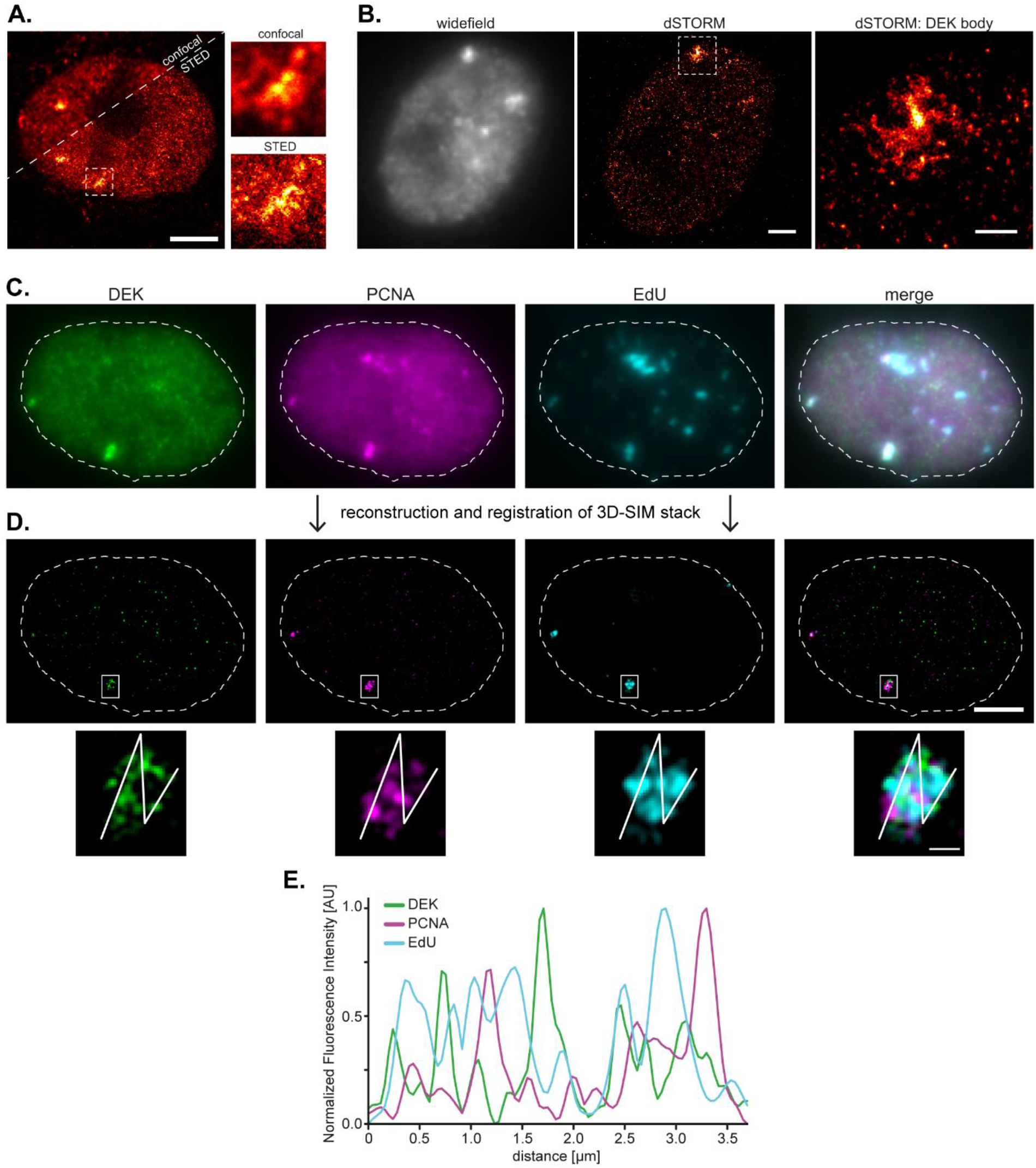
DEK bodies localize in the vicinity of the replication fork. **A.** Confocal and STED super-resolution images of an MCF10A cell nucleus in late S- phase labeled with anti-DEK antibodies. The STED image reveals a granular internal structure of DEK bodies (insets). Scale bar: 5 μm. **B.** dSTORM images of an MCF10A cell nucleus in late S-phase labeled with anti-DEK antibodies. Left: widefield conventional image of DEK immunolabelling. Middle: dSTORM image reconstructed from single-molecule localization microscopy acquisition. Scale bar: 2 μm. Right: magnified image of the ROI shown in the middle containing a DEK body. Scale bar: 0.5 μm. **C.** Pseudo-widefield representation of a representative U2-OS nucleus in late S-phase imaged in 3D-SIM mode. U2-OS cells were pulse-labelled with EdU for 10 min. DEK (green) and PCNA (magenta) were visualized *via* indirect immunofluorescence microscopy and EdU (cyan) using click chemistry. **D.** Single Z-slice from a middle section of the super-resolved image stack after reconstruction and registration. Scale bar: 5 μm. The magnified inset shows one DEK- positive large replication focus. Scale bar: 0.5 μm. **E.** Fluorescence intensity profiles of DEK, PCNA and EdU. Interpolated intensity profiles were calculated along the line shown in (D) and normalized to the min/max values.

### DEK bodies assemble at replicating heterochromatic regions

The occurrence of DEK in a subpopulation of PCNA- and EdU-positive nuclear bodies in late S-phase let us hypothesize that DEK accumulates at sites of replicating heterochromatin. To test this hypothesis, we conducted immunofluorescence analyses of three well-established chromatin markers in S-phase cells - H3K9ac for euchromatin (Wang et al., 2008), H3K27me3 for facultative heterochromatin (Trojer and Reinberg, 2007), and H3K9me3 for constitutive heterochromatin (Nakayama et al., 2001) (**Fig. 3A, B** and **Fig. S3 A-C**). We observed that within DEK bodies, DEK-specific signals did not overlap with H3K9ac labeled chromatin, were spatially juxtaposed to H3K27me3 signals, and colocalized with H3K9me3 (**Fig. 3A**), strongly suggesting that DEK bodies represent regions of replicating constitutive heterochromatin. This conclusion was corroborated by Manders’ coefficient analysis yielding the highest overlap for DEK and H3K9me3. The values for the coefficient M_1_ indicating the fraction of DEK bodies containing the different modified histones were as follows: M_1(DEK&H3K9me3)_ = 0.87 ± 0.09, M_1(DEK&H3K9ac)_ = 0.22 ± 0.12; and M_1(DEK&H3K27me3)_ = 0.37 ± 0.06; with n=8 from two independent experiments (**Fig. 3B**). Further evidence was obtained by Proximity Ligation Assay (PLA) confirming direct interaction of DEK with H3K9me3 but not with H3K27me3 (**Fig. 3C, D**), while fluorescence recovery after photobleaching (FRAP) showed a reduced mobility of DEK in the bodies with respect to the bulk chromatin, consistent with a high level of chromatin compaction in these structures (**Fig. 3E-G**). In line with these data, DEK bodies were associated with heterochromatin, in particular with H3K9me3, also in U2-OS wild type and U2-OS KI eGFP-DEK cells (Fig. S4).

**Figure 3:**
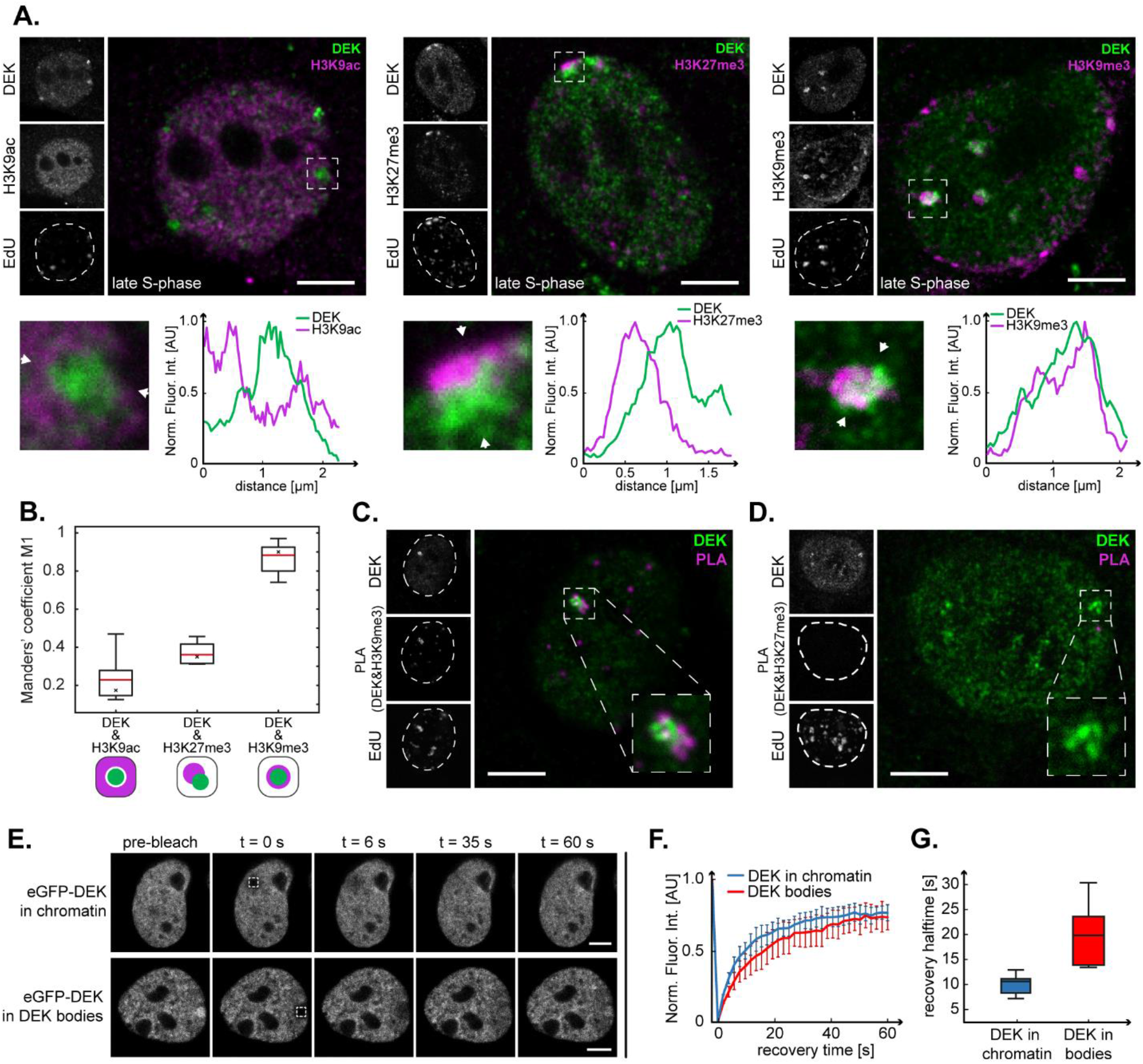
DEK bodies position within constitutive heterochromatin regions in late S-phase. **A.** Confocal images of late S-phase MCF10A cell nuclei labelled with EdU and antibodies specific for DEK and for three post-translational histone modifications (H3K9ac - left, H3K27me3 - centre, H3K9me3 - right). ROIs containing DEK bodies are marked by a dashed line and displayed as magnified images below. The fluorescence intensity profiles were calculated along the lines delimited by the white arrowheads and normalized to 1. Scale bars: 5 μm. **B.** Analysis of colocalization of DEK bodies and chromatin marks using Manders’ coefficients. In the box plots, the red line indicates the average value, the asterisk marks the median, the box boundaries represent the interquartile range, and the whiskers above and below the box indicate minimum and maximum values. Graphs on the bottom schematically represent localization pattern. **C.** Confocal image of a late S-phase MCF10A cell nucleus after PLA with antibodies specific for DEK and H3K9me3. The PLA positive signal (magenta) indicates direct interaction. Scale bar: 5 μm. **D.** Confocal image of a late S-phase MCF10A cell nucleus after PLA with antibodies specific for DEK and H3K27me3. No PLA positive signal is detected. Scale bar: 5 μm. **E.** FRAP analysis of DEK-eGFP mobility transiently expressed in MCF10A cells in bulk chromatin and in DEK bodies. Bleach-ROIs (2 μm x 2 μm) are indicated by dashed squares. Fluorescence recovery was monitored for 3 min. Scale bar: 5 μm. **F.** DEK displays a lower mobility in bodies than in bulk chromatin. Normalized FRAP recovery curves based on the data from (E). Data points show the average of 6 experiments for DEK bodies and 14 experiments for DEK in chromatin. Error bars show the SEM. **G.** Halftimes of recovery for eGFP-DEK in chromatin (τ½ = 10.6 s ± 3.1 s), and in DEK bodies (τ½ = 17.1 s ± 7 s) calculated from the FRAP data in (F).

Finally, we investigated the localization of DEK bodies with respect to known heterochromatic compartments such as centromeres, the Barr body, and the nuclear periphery. Maximum intensity projections of confocal Z-stacks revealed that the epigenetic mark for centromeres, the histone H3 variant CENP-A, reproducibly decorates DEK bodies from both sides (**Fig. 4A**). We did not observe spatial proximity of DEK and CENP-A outside of late S (**Fig. 4B**). A staining pattern of similar symmetry was observed for *Xist* RNA, a selective marker for the Barr body (Cerase et al., 2015) (**Fig. 4C**). No such pattern was observed in male BPH-1 cells (data not shown). Similarly, we found DEK bodies in close proximity of the nuclear periphery and Lamin-A positive nuclear invaginations only in S-phase **(Fig. 4D, E**). Furthermore, DEK bodies corresponded to regions of dense chromatin, as shown by DNA staining with ToPro3 (**Fig. 4F**). Altogether, our collective microscopy data strongly support a role for DEK bodies in the replication of heterochromatic chromatin regions in late S-phase.

**Figure 4:**
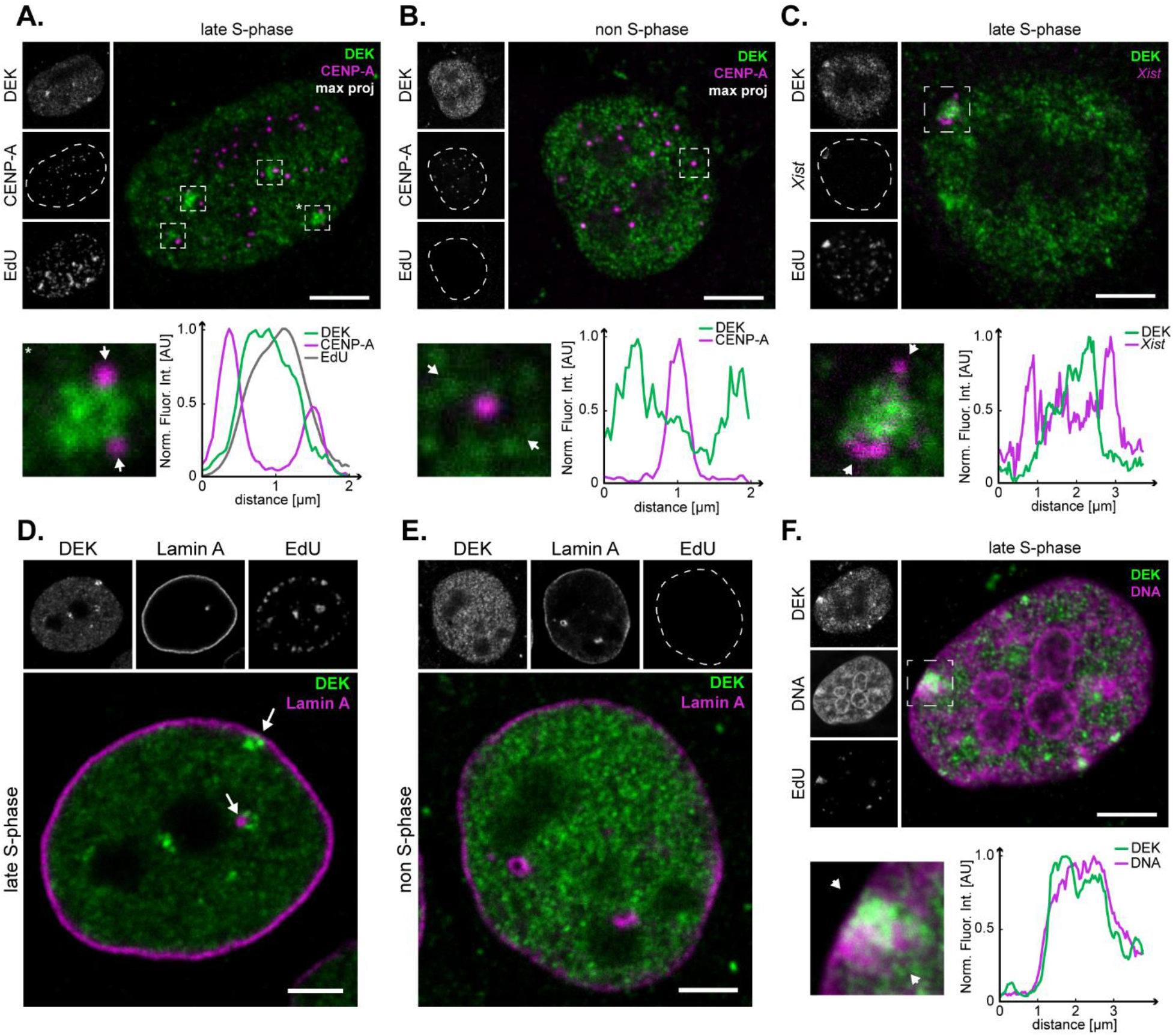
DEK bodies are spatially associated with replicating heterochromatin. **A, B**. Maximum intensity projection images of representative MCF10A cell nuclei in late S-phase (A), and in the absence of DNA replication (B, non-S-phase, EdU-negative) labelled with anti-DEK, anti-CENP-A antibodies, and EdU-AF467. ROIs are marked by a dashed line and displayed as magnified images below. The fluorescence intensity profiles were calculated along the lines delimited by the white arrowheads and normalized between 0 and 1. Scale bars: 5 μm. **C**. Maximum intensity projection image of a representative MCF10A cell nucleus in late S-phase labelled with anti-DEK antibodies, a *Xist* RNA-specific FISH probe, and EdU- AF467. The ROI marked by a dashed line is displayed as magnified image below. The fluorescence intensity profile was calculated along the lines delimited by the white arrowheads and normalized between 0 and 1. Scale bars: 5 μm. **D, E**. Confocal images of representative MCF10A cell nuclei in late S-phase (D) and in the absence of replication (E, non-S-phase, EdU-negative) labelled with anti-DEK and anti-Lamin A antibodies, and EdU-AF467. Arrows in (D) indicate the juxtaposition of DEK and Lamin A at the periphery of the cell nucleus and at nuclear lamina invaginations, which is not observed in non-S-phase cells (E). Scale bar: 5 μm **F.** Confocal image of a representative MCF10A cell nucleus in late S-phase labeled with DEK-specific antibodies, with the DNA dye ToPRo3 and EdU-AF647. The ROI marked by a dashed line is displayed as magnified image below and shows a DEK body overlapping with a region of dense chromatin. The fluorescence intensity profile was calculated along the lines delimited by the white arrowheads and normalized between 0 and 1. Scale bars: 5 μm.

### DEK body dynamics is affected by DNA replication inhibitors

Having assessed that DEK bodies are implicated in the replication of heterochromatin in fixed cells, we were interested in exploring and manipulating their dynamics in living cells. To this end, we transiently expressed mRFP-PCNA in U2-OS KI eGFP-DEK cells *via* a PCNA chromobody plasmid and performed time-lapse imaging experiments at a spinning disk confocal microscope (**Fig. 5A**). DEK body formation was evaluated manually and occurred reproducibly in every cell in late S-phase. 2.7 ± 1 DEK bodies formed per cell and persisted for 48 ± 30 min until G2 entry (**Fig. 5B, C**). All DEK bodies colocalized with PCNA-positive replication foci, but only 45% of the latter contained DEK. A similar behavior was observed also in MCF10A cells transiently expressing fluorescent fusions of DEK and PCNA, with a higher number of DEK bodies (6.7 ± 2.2) that persisted for a longer time (189 ± 38 min). 82 ± 18% of DEK bodies colocalized with PCNA-marked replication foci in these cells (object-based analysis of 19 DEK bodies) (**Fig. S5**).

**Figure 5:**
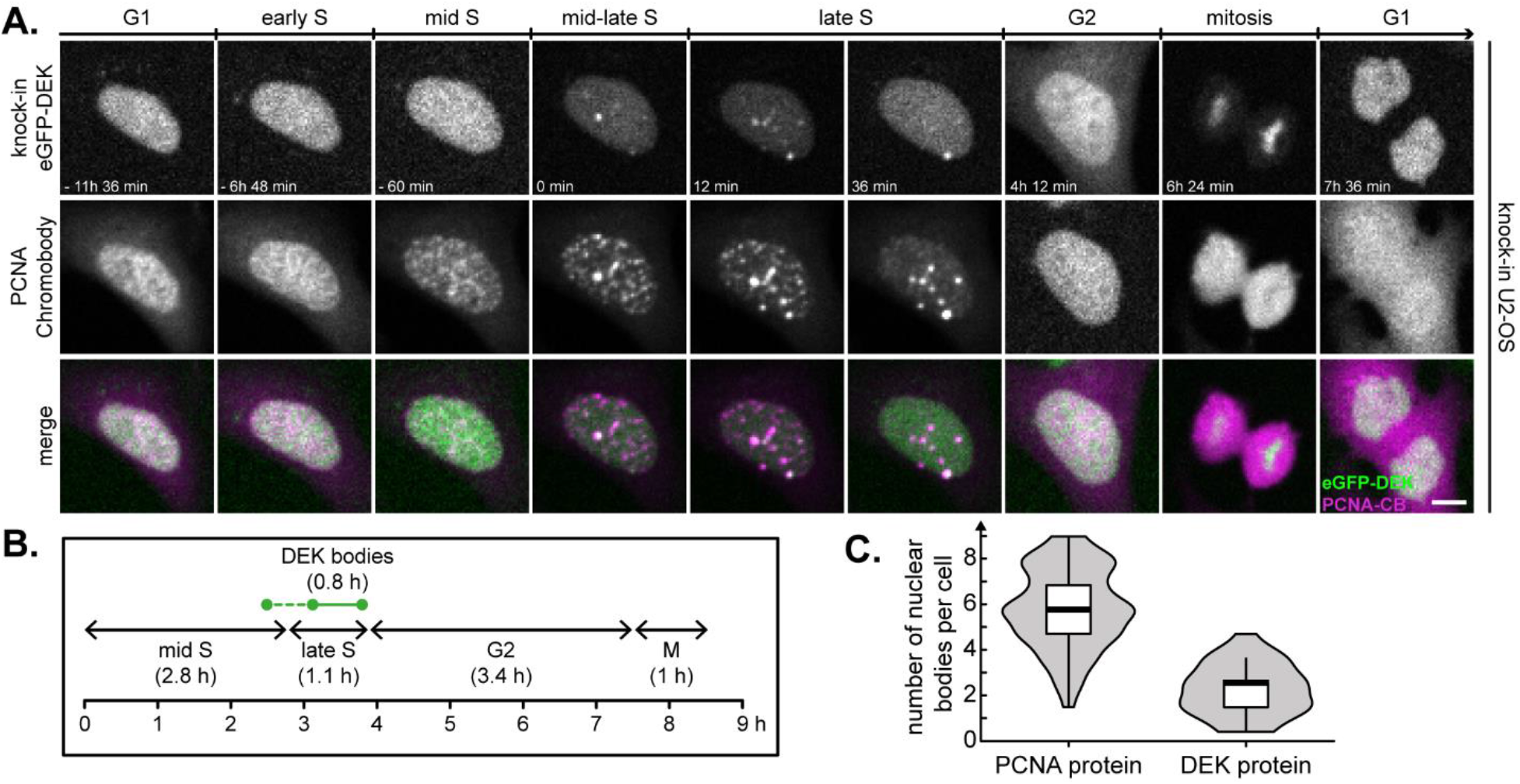
DEK bodies form during late S-phase of the cell cycle. **A.** Time-lapse fluorescence microscopy of U2-OS KI eGFP-DEK cells transiently expressing mRFP-PCNA. Images were acquired at a spinning disk confocal microscope. The figure shows a representative image sequence. Magenta: PCNA chromobody, green: eGFP-DEK. Scale bar: 5 μm. **B.** Mean durations of cell cycle phases and mean DEK body lifetime. A total of 56 cells were scored manually from two independent experiments. Mid S, late S, G2 and M-phases were identified according to the PCNA localization pattern. Only images with clearly detectable DEK bodies were scored as positive. DEK bodies appear reproducibly in late S-phase (continuous green line), and sporadically also in mid-S-phase (dashed green line). **C.** Quantification of PCNA body and DEK body numbers in late S-phase using the same data set as in (B). Only PCNA or DEK bodies were included in the evaluation that persisted for at least two consecutive frames. The violin plot shows the density distribution of the data, the black line indicates the median, the box the interquartile range (IQR) and the whiskers the IQR ± 1.5.

To determine whether impairment of DNA replication would affect DEK body formation, we employed hydroxyurea (10 μM), aphidicolin (200 nM) and camptothecin (100 nM), three well-established replication fork inhibitors acting by different mechanisms which at these low doses slow down but do not arrest replication (**Fig. 6A**) (Deutzmann et al., 2015; Vesela et al., 2017). The treatments, in particular aphidicolin and camptothecin, did not affect viability, but triggered accumulation of cells in G2, followed by normal mitoses (**Fig. 6B upper panels**). Of the three substances, aphidicolin showed the most prominent effect reducing DEK body number by 47%, and prolonging DEK body lifetime by 28%. Camptothecin had a similar but more moderate effect, and hydroxyurea did not alter DEK body dynamics (**Fig.6B lower panels**). The differential response to the replication stress inducers strongly suggests that DEK bodies are implicated in the replication of DNA structures particularly susceptible to aphidicolin, such as common fragile sites that in turn can coincide with highly repetitive heterochromatic regions like centromeres or ribosomal DNA (Lukusa and Fryns, 2008), in line with our data on the colocalization of DEK bodies with heterochromatin histone marks.

**Figure 6:**
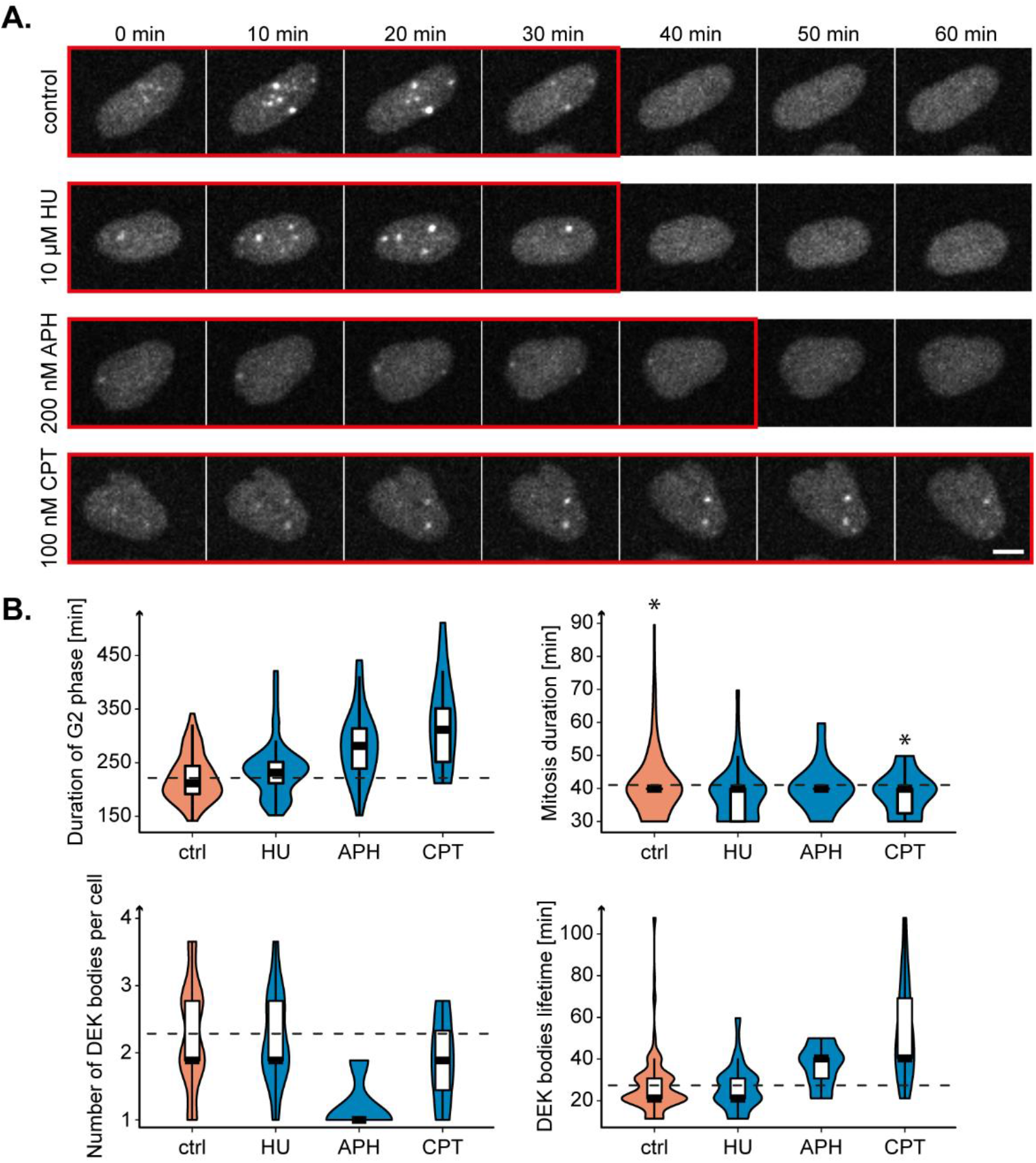
Formation of DEK bodies is dependent on DNA replication. **A.** Time-lapse fluorescence microscopy of U2-OS KI eGFP-DEK cells treated for 1 h with either 10 μM HU, or 200 nM APH, or 100 nM CPT or left untreated (Ctrl). Images were acquired for 24 h with a spinning disk confocal microscope. A representative time series is shown for each treatment condition. Frames with DEK bodies are highlighted by a red outline. Scale bar: 10 μm. **B.** Quantification of G2-phase and mitosis duration, DEK body number and lifetime in time series data as in (A) (15-132 cells, single experiment). The violin plots show the density distribution of the data points, the black line indicates the median, the box the interquartile range (IQR) and the whiskers the IQR ± 1.5. (*) The control and CPT data sets contained one outlier each, which was omitted from the quantification.

### Down-regulation of the SUMO pathway increases DEK body number in an siRNA screen for DEK body regulators

As our time-lapse microscopy experiments could measure the effect of mild replication stress on DEK body formation, we chose to perform an imaging-based unbiased approach to identify DEK body regulatory factors. To this end, we set up a high-throughput siRNA screen targeting genes from a recently published DEK interactome (Smith et al., 2018), DEK-related factors derived from the literature, and genes selected from a commercial library targeting the DNA damage response that responded to the gene ontology terms “DNA replication” and “S-phase”. The final library consisted of 332 genes (678 siRNAs total, at least two siRNAs per gene, **Supplementary Table S1**).

Experimental conditions and a suitable image analysis pipeline using CellProfiler software were established in two pilot screens including aphidicolin and camptothecin as positive controls (**Supplementary Material and Methods**, **Supplementary Table S2** and **Fig. S6**). The primary screen targeting the chosen gene panel was performed in duplicate and a Z-score of the log-transformed mean number of DEK bodies per cell was calculated for each siRNA (**Fig. 7A**). This resulted in 51 candidate siRNAs enhancing (up-regulators) and 29 candidates reducing DEK body numbers (down-regulators) corresponding to 44 and 28 different genes, respectively (**Fig. 7B, C**). The number of DEK bodies per cell was the most sensitive and robust readout for DEK body formation also in the automated approach, reproducing the results from the manual experiments (**Fig. S6D**). To further confirm the identified hits, we conducted a low-throughput secondary screen with 28 siRNAs (**Fig. 7D**). In general, the values for the Z-score obtained in this secondary screen were smaller and more variable, most likely due to the different, less automated imaging system and the manual dispensing of transfection reagents. Still, the effect of up and down-regulation of DEK body number was confirmed (**Fig. 7D**). Importantly the secondary screen validated four top DEK body up-regulators from the primary screen belonging to the SUMO pathway, among them the E1 activating enzyme SAE1 and the E2 conjugating enzyme UBE2I. Our screening approach thus determined that the SUMO pathway plays a major role in the regulation of DEK body formation.

**Figure 7:**
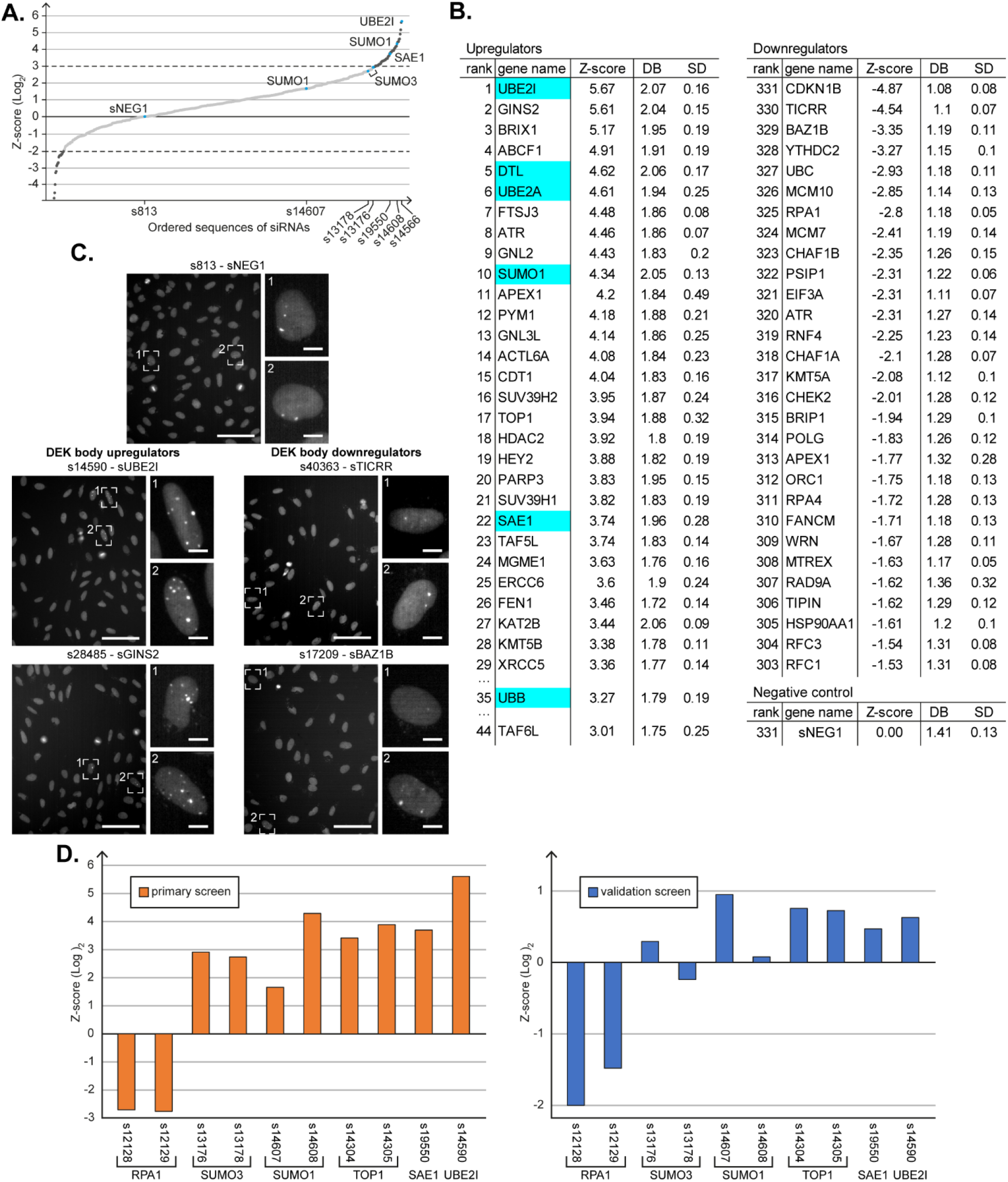
siRNA screen identifies positive and negative regulators of DEK body formation. **A.** Plot of the log2-transformed ranked mean Z-score data of the primary screen. The screen was performed on 4 field of view per well, with two wells per siRNA (for each 2 replicates), yielding 16 Z-scores for each siRNA. Dark grey dots: Top scorer (3 ≤ Z-score ≤ -2). Blue dots: Target genes included in the pilot screen. sNEG1: negative control siRNA. **B.** List of the strongest DEK body upregulating and downregulating genes based on the Z-score of the corresponding siRNAs. For the up-regulators, ranks 30-34 and 36-43 are not shown for better readability. Mean number of DEK bodies (DB) and standard deviation (SD) are shown. Highlighted are genes belonging to the SUMO pathway. **C.** Representative widefield images from the primary screen showing U2-OS KI eGFP-DEK transfected with a negative control siRNA sNEG1 (s813) and siRNAs targeting the two strongest DEK body upregulating (left) and downregulating (right) genes. Magnified insets with DEK body-positive nuclei are shown. Scale bars: 100 μm; inset scale bars: 10. μm **D.** Comparison of z-score data from the primary and the secondary (validation) screen. The log2-transformed mean Z-score for the indicated siRNAs is shown. The primary screen was performed in duplicate, the validation screen in triplicate. Knockdown efficiency of SUMO1 and SUMO 3 siRNAs was confirmed by Western Blot (Fig. S7)

### DEK is a SUMO target and SUMOylation is required for the formation of DEK bodies

As both screening approaches showed that down-regulation of the SUMOylation pathway resulted in the up-regulation of the number of DEK bodies in U2-OS KI eGFP-DEK cells, we hypothesized that DEK may be a target of SUMOylation and that, in turn, its SUMOylation state may modulate the ability to form DEK bodies in late S-phase. Indeed, the DEK protein sequence contains a SUMOylation consensus motif ψ-K-X-E/D (Zhao et al., 2014) at lysine 261 (AKRE, https://sumo.biocuckoo.cn/online.php) and DEK has been shown to be SUMOylated under stress conditions in a mass spectrometry study of H_2_0_2_- treated SH-SY5Y neuroblastoma cells (Grant, 2010). To test if DEK can be SUMOylated under steady-state conditions, we focused on SUMO 1 and 3, as both were strong hits in the screens. We first conducted co-immunoprecipitations using DEK-antibodies, showing direct interaction of DEK with SUMO1 and SUMO2/3 (**Fig. 8A**). To narrow down a specific SUMO paralog targeting DEK, we used HeLa cells stably expressing 6xHis-SUMO-1, 6xHis-SUMO-2, or 6xHis-SUMO-3 allowing for subsequent nickel affinity purification of SUMOylated factors. This approach revealed modification of DEK by all SUMOs, yet with the strongest activity for SUMO-3 (which also generates multimers) (**Fig. 8B, C**). As we could confirm the existence of SUMOylated DEK in cells under physiological conditions, we were next interested to map the specific SUMOylation sites in the DEK primary sequence. As indicated above, DEK harbors one SUMOylation consensus motif at position lysine 261. However, mutation of this site did not yield substantial reduction in SUMOylation in *in vitro* SUMOylation assays using recombinant GST-tagged DEK as acceptor (**Fig. S8**). Therefore, we screened the DEK sequence for additional potential SUMOylation sites, which revealed amino acids 61, 62 and 144, 145 as suitable SUMO attachment sites. Interestingly, mutation of these sites individually, as was the case with 261, did not result in a substantial reduction of SUMOylation activity on DEK (**Fig. S8**). However, DEK molecules carrying mutations at all five sites showed an 80% reduction in SUMOylation (**Fig. 8D, E**), highlighting that most of the potential SUMOylation sites were captured by our mutational approach. Given that we had created a DEK mutant that is substantially less susceptible to SUMOylation, we then investigated the ability of such a mutant to form DEK bodies in cells. Therefore, we established a stable cell line expressing this mutant as eGFP-fusion and monitored the occurrence of DEK bodies as described above. No DEK body formation was observed in cells expressing SUMO-less DEK (GST DEK SUMO mut, **Fig. 8F**). This result points at a complex regulation of DEK bodies by SUMO, where SUMOylation of DEK seems to be required for DEK body formation while DEK body numbers increase when the overall SUMOylation capacity of the cell is reduced. In summary, these data uncover a yet unrecognized link between SUMO and DEK and establish SUMOylation as a crucial regulator of DEK body formation.

**Figure 8:**
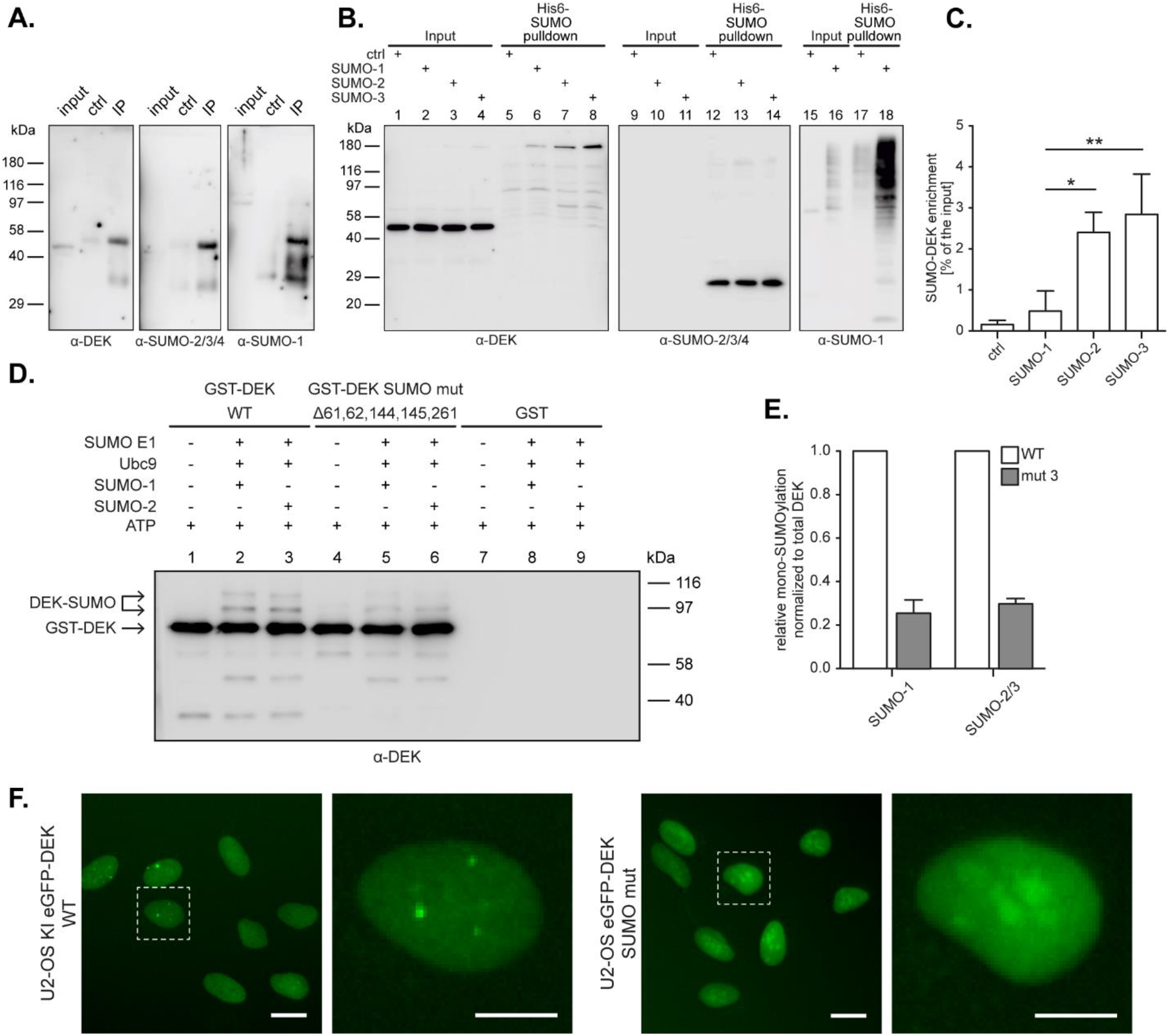
DEK is SUMOylated in cells and SUMOylation is involved in the formation of DEK bodies. **A.** Immunoblot analysis of co-immunoprecipitations from U2-OS whole cell lysates using DEK-, and SUMO-specific antibodies. **B.** Immunoblot analysis of His-SUMO pulldown experiments from HeLa cells expressing 6xHis-SUMO-1, 6xHis-SUMO-2, 6xHis-SUMO-3, or control cells after nickel affinity chromatography. Purified proteins were analyzed with DEK and SUMO-specific antibodies. **C.** Densitometric quantification of mono-SUMOylated DEK signals from (B, left panel). DEK signals in pulldown samples were normalized to the corresponding input samples. Error bars indicate the SD. **p<0.01, *p<0.05. **D.** *In vitro* SUMOylation assay: Immunoblot analysis of *in vitro* SUMOylation reactions containing recombinant GST-tagged DEK WT, mutant, or GST only proteins as indicated and using DEK-specific antibodies for detection. **E.** Densitometric quantification of mono-SUMOylated DEK from *in vitro* samples as in(D). Error bars indicate the SD. **F.** Widefield microscopy images of U2-OS KI eGFP-DEK or a cell line stably expressing a wild-type or a SUMOylation deficient DEK mutant - DEKmut (Δ61,62,144,145,261) cells. Scale bars: 20 μm.

## Discussion

### DEK bodies reflect the formation of transient DEK multimers in the nucleus during late DNA replication

We report on the discovery of DEK bodies, DNA replication-dependent focal assemblies of the oncoprotein DEK in the cell nucleus. DEK bodies were observed in every cell during mid-late S-phase in primary, immortalized and in transformed, yet non-malignant cell lines. This finding was somewhat surprising as the large majority of immunofluorescence studies have described DEK as a factor dispersed in an uniform punctuate pattern in the nucleus (Hu et al., 2007; Kappes et al., 2001; Matrka et al., 2015). Intriguingly, the existence of non-covalent DEK multimers was proposed in the past based on biochemical analyses of DEK-chromatin complexes (Kappes et al., 2004b), but had eluded demonstration in live cells so far. We interpret DEK bodies as the cellular correlate of these non-covalent multimers, whose detection by microscopy was most likely facilitated by fusion of DEK to GFP. This reporter has a propensity to form aggregates at high concentrations (Stepanenko et al., 2021) contributing to slow down the dynamics of DEK bodies and make them amenable to microscopic imaging. The focused inspection of high resolution immunofluorescence images revealed that DEK bodies are distinct sites of active replication containing endogenous DEK in close spatial proximity to nascent DNA and the replication processivity factor PCNA consistent with previous iPOND analyses (Alabert et al., 2014; Aranda et al., 2014; Garcia et al., 2017; Lossaint et al., 2013; Ribeyre et al., 2016; Sirbu et al., 2013). The number and persistence of DEK bodies varied substantially between MCF10-A and U2-OS cells, the former showing a more pronounced expression of these bodies. Unfortunately, a direct quantitative comparison of DEK bodies between these two cell lines is not appropriate as DEK was expressed endogenously in U2-OS and transiently in MFC-10A cells, and imaging was performed with different microscopes in different laboratories. But in both cases, DEK bodies were sparingly observed also in mid S-phase, probably reflecting cell-to-cell variability in replication timing (Zhang et al., 2017).

Previous data suggesting that DEK may occur in defined nuclear substructures included coimmunoprecipitation and spurious colocalization with PML (Ivanauskiene et al., 2014), and the redistribution to interchromatin granule clusters in the presence of deacetylase inhibitors (Clarey 2005, McGarvey 2000), but no link to the cell cycle or DNA replication was made in these studies.

In general, the formation of nuclear bodies has emerged as an organizing principle for the orchestration of genome-associated functions in time and space. Different nuclear bodies have been described in which proteins, often together with RNAs, are sequestered and released according to demand, including the nucleolus, nuclear speckles and paraspeckles, PML, Cajal bodies, 53BP1 and other bodies (Dundr, 2012). Post translational modifications, as well as weak intermolecular interactions mediated by intrinsically disordered protein domains, seem to play a fundamental role in the genesis and dynamic composition of these membrane-less nuclear condensates. It is thus not surprising that DEK, a DNA and RNA-binding protein carrying several post-translational modifications and an intrinsically disordered region (aa 170-270) is prone to form nuclear bodies.

### DEK bodies form at heterochromatic regions

Our data provide strong evidence that DEK bodies are sites of heterochromatin replication. These include their narrow window of appearance during late S-phase, the association with heterochromatin histone marker H3K9me3, the high level of chromatin compaction and the reduced mobility of DEK in bodies, and the local proximity to the Barr body and centromeric chromatin CENP-A. The lack of colocalization between CENP-A and DEK reported previously (Ivanauskiene et al., 2014) was most likely due to the fact that endogenous DEK bodies are transient structures visible only in late S-phase, and may thus escaped detection.

The reproducibility with which DEK bodies form in consecutive cell cycles support the notion that they reflect a conserved and necessary passage in genome duplication. The striking appearance of the centromeric and *Xist*-specific signals as dots decorating DEK bodies (Fig. 4A, C) are suggestive of newly replicated chromosomes tethered to these bodies at regions rich of secondary structures that are preferred DEK substrates *in vitro*, are difficult to replicate and prone to breakage. In line with this observation, downregulation of DEK resulted in the formation of anaphase bridges, the appearance of DNA breaks on mitotic chromosomes and of 53BP1 bodies in daughter cells (Deutzmann et al., 2015). While, in the latter study we described how DEK affects DNA replication globally, the results obtained here suggest a more specialized function of the protein, at specific genomic regions including the centromere and possibly other constitutive heterochromatic regions. These findings may explain the relatively mild effects of DEK downregulation on replication fork progression as assessed by bulk fiber assays. The assessment of DEK′s influence on the processivity of the replication fork therefore awaits DNA combing experiments coupled with the detection of heterochromatic fibers.

### DEK bodies are novel players in the SUMO-dependent regulation of heterochromatin replication

DEK body up-regulating siRNAs show a remarkable enrichment in proteins involved in post-translational modification pathways. Histone acetyltransferases (HATs) ACTL6A, KAT2B, TAF5L and TAF6L, histone deacetylase HDAC2, histone methyltransferases SUV39H1/2 and SUMOylation-related proteins UBE2I, SUMO1 and SAE1 are among the top hits. Acetylation of DEK by KAT2B/PCAF has been shown previously (Cleary et al., 2005). In this study, overexpression of KAT2B led to an accumulation of DEK in interchromatin granule clusters (IGCs), which are nuclear domains enriched in pre-mRNA splicing factors. Pharmacological inhibition of KAT2B in turn, blocked the relocation of DEK into IGCs, opposite to the effect of downregulation of KAT2B on DEK bodies. This finding further underscores that DEK bodies are distinct from ICGs, which are more abundant (20-30 clusters per cell) and are stable during the whole interphase (Mintz et al., 1999).

Here, we focused on SUMOylation as a post-translational modification that has recently emerged as a major regulator of heterochromatin formation and replication. We substantiate the result from the siRNA screen by showing that DEK is modified by SUMO in cells, and that SUMO-less DEK does not form bodies. Recent data also show that SUV39H1 promotes SUMOylation and *de novo* targeting of HP1α to pericentric heterochromatin, independently of its methyltransferase activity. The latter is required in a second step to maintain and propagate heterochromatin (Maison et al., 2016). This knowledge aligns well with our screening data, where downregulation of both SUV39H1 and SUMO increase the number of DEK bodies suggesting the DEK plays a role in “seeding” constitutive heterochromatin after DNA replication (Maison et al., 2016). Our data also support and refine the conclusions from our previous study showing that DEK is essential for heterochromatin integrity, and has a Su(var)like activity by enhancing HP1α binding to H3K9me3 (Kappes et al., 2011a).

Heterochromatin preferentially harbors repetitive sequences causing non-B DNA secondary structures. Their replication requires specialized polymerases that can sustain fork progression in the presence of difficult-to-replicate regions, but on the other hand lack proofreading activity and are therefore error-prone and mutagenic (reviewed in (Tsao and Eckert, 2018)). The switch from replicative polymerase to these specialized, or translesion polymerases, at the replisome is regulated by ubiquitylation and SUMOylation (Despras et al., 2016; Mailand et al., 2013). In particular, SUMOylation is required on the one hand, for driving specialized polymerases to nascent DNA while, on the other hand, enabling discharge from the replisome after bypass of the problematic site to limit replication errors and prevent mutagenesis, as shown for Pol η (Despras et al., 2016; Guérillon et al., 2020). We hypothesize first, that DEK in bodies may participate in the maturation of nascent heterochromatin after replication involving specialized polymerases and that this step requires SUMOylation of DEK, explaining why a non-SUMOylatable mutant of DEK does not form bodies. Secondly, we interpret the increase in the number of DEK bodies observed after downregulation of the SUMO pathway as consequence of impaired turnover of these specialized replisomes and the replacement by proof-reading replicative polymerases. Interestingly, also Polη was reported to be SUMOylated at multiple sites which can substitute for each other, so that simultaneous mutation of all SUMOylation sites was reported to be necessary to achieve dysregulation of the Polη dynamics (Guérillon et al., 2020).

At present, we have no straightforward interpretation for the reduction in DEK bodies that seems to result from increased DNA replication stress, as highlighted by the lack of DEK bodies in highly cancerous MDA-MB-231 cells and in the presence of aphidicolin, and - to a lesser extent - camptothecin. We speculate that under these conditions replisomes at difficult-to-replicate-regions would be more likely to collapse leading to DNA breakage before chromatin maturation, thus impairing DEK body assembly. Accordingly, we found increased γH2AX foci after DEK downregulation in a previous study (Kappes et al., 2008). The propensity to DNA strand break formation is a well-established feature of difficult-to-replicate-regions such a common fragile sites (CFS) and their stability has been shown to depend on specialized polymerases (Bergoglio et al., 2013; Bhat et al., 2013).

In conclusion, the link established here between DEK and the establishment of constitutive heterochromatin during replication sheds new light on the mechanisms of action of this so far enigmatic oncoprotein. Some of its pleiotropic effects may be associated with the seeding of (constitutive) heterochromatin, a process that broadly impacts the celĺs epigenetic landscape and transcriptional program and is essential for maintaining genomic stability (Penagos-Puig and Furlan-Magaril, 2020).

## Materials and Methods

### Cell culture

Mammary epithelial, non-transformed cell line MCF10A (ATCC CRL-10317) was grown in DMEM:F-12 (Dulbecco’s Modified Eagle Medium:Nutrient Mixture F-12) (1 : 1) medium (Gibco, 11330057) supplemented with 5% horse serum (HS), 2 mM L-glutamine and 1% penicillin/streptomycin (Sigma-Aldrich, G6784), 10 μg/ml insulin (Sigma-Aldrich, I9278), 0,5 μg/ml hydrocortisone (Sigma-Aldrich, H0888), and 20 ng/ml of human Epidermal Growth Factor (hEGF, Sigma-Aldrich, E9644) added freshly before the use of full medium. MCF7 cells and MDA-MB-231 cells (ATTC HTB-26) were grown in DMEM medium (Gibco, 11330057) supplemented with 10% fetal bovine serum (FBS, Euroclone, ECS0180L), 1% penicillin/streptomycin (Sigma-Aldrich, G6784) and 2 mM L-glutamine. Cells were grown on 10 cm^2^ dishes (about 10-12 cell passages) at 37°C in 5% CO_2_. For any experiments on fixed cells, cells were plated on glass coverslips coated with 0.5% (w/v) pork gelatine (Sigma-Aldrich, G2500) dissolved in phosphate buffer saline (PBS) and autoclaved. U2-OS osteosarcoma (kind gift from G. Marra, University of Zurich, Switzerland) and U2-OS KI eGFP-DEK cells (described in (Ganz et al., 2019)) were cultured in McCoy’s 5a modified medium (Gibco, Life Technologies) supplemented with 10 % FBS (Capricorn Scientific), 100 U/ml penicillin and 100 μg/ml streptomycin (both Gibco, Life Technologies). For selection after transfection, Geneticin (Thermo Fisher Scientific) was added at concentrations of 200 μg/ml for U2OS KI eGFP-DEK WT, eGFP-DEK SUMOmut Δ61,62, eGFP-DEK SUMOmut Δ261 and eGFP-DEK SUMOmut Δ61,62,144,145,261 cells. During all experiments using siRNA-mediated protein down regulation penicillin and streptomycin were omitted from the medium. HeLa S3 cervical adenocarcinoma cells and HeLa 6xHis-SUMO cells (kind gift from Ronald T. Hay, University of Dundee, UK) were cultured in DMEM medium (Gibco, Life Technologies) supplemented with 10% FBS (Capricorn Scientific), 100 U/ml penicillin, 100 μg/ml streptomycin (both Gibco, Life Technologies) and 6 mM L-glutamine (Gibco, Life Technologies). hTERT immortalized BJ-5ta foreskin fibroblasts were cultured in a 4:1 mixture of DMEM medium and Medium 199 (Gibco, Life Technologies) supplemented with 10% FBS, 4 mM L-glutamine and 10 μg/ml hygromycin B (Calbiochem).

### Transfection and cell sorting

For detection of endogenous PCNA, U2-OS KI eGFP-DEK or MCF10A cells were transfected with a PCNA-specific chromobody expression vector (pCCC-TagRFP, Chromotek) using Effectene (Qiagen U2-OS KI eGFP-DEK) or Lipofectamine 3000 (Invitrogen) according to the manufactureŕs instructions. For U2-OS KI eGFP-DEK, the medium was exchanged 4 h after transfection and the cells were incubated for further 20 h before imaging. For MCF10A cells, the medium was exchanged about 12 h post transfection, and experiments were carried out between 24 to 48 h after transfection. For the generation of U2-OS cell lines stably expressing eGFP-DEK mutants, pEGFP-N1- hDEK plasmids encoding the mutated DEK versions were transfected into U2-OS DEK knockout cells using Lipofectamine 3000 reagent (Invitrogen). 3-9 days after transfection (depending on cell growth and viability) selection was carried out with geneticin at a final concentration of 200 μg/ml. To create cell lines with a uniform GFP fluorescent signal, the stable transfected cell lines were bulk sorted for a moderate fluorescence signal using a FACSAria IIIu Cell Sorter (BD Biosciences).

### Immunochemical methods

All antibodies used in this study are listed in **Supplementary Table 3**.

For the detection of nascent DNA MCF10A and U2-OS cells were incubated with EdU (10 μM) for 25 min or 10 min respectively. After fixation and washings, the incorporated precursor was detected with the Click-iT EdU Imaging Kit or Click-iT EdU PLUS Imaging Kit, Molecular Probes with Alexa Fluor 647 or Alexa Fluor 594 azide dye. When necessary, nuclei were counterstained with TO-PRO™-3 Iodide (Invitrogen, 1:2000). For immunofluorescence detection of DEK or PCNA cells were fixed with 4% PFA/PBS (20 min) followed by quenching with 50 mM NH_4_Cl/PBS (10 min). Cells were permeabilized in methanol (5 min, -20°C), incubated in 1% BSA/PBS for 30 min followed by incubation with the primary antibody dilution (diluent 1% BSA/PBS) in a dark, humid chamber (overnight, 4 °C). After washing with PBS for three times, cells were incubated with the appropriate secondary antibody (diluent 1% BSA/PBS) in the dark chamber (1 h). Coverslips were mounted either in ProLong™ Diamond Antifade Mountant (Invitrogen, P36961) or Aqua Polymount (Polysciences, 186065).

### RNA-FISH

*Xist* RNA was detected with Stellaris® RNA FISH (Human XIST with Quasar® 570 Dye, Biosearch Technologies, SMF-2038-1) according to the manufacturer’s instructions. After the hybridization step, Click-iT EdU detection was performed with the use of Alexa Fluor 647 azide. The samples were washed with PBS and mounted in ProLong™ Diamond Antifade Mountant.

### Proximity Ligation Assay

Proximity Ligation Assay (PLA) was performed in MCF10A cells using the DuoLink® in situ Orange detection reagent (DUO92102, Sigma-Aldrich) according to the manufacturer’s instructions. Cells were incubated for 60 min at 37°C in a humidified chamber in PLA blocking solution, followed by 1h at RT with antibody solution containing primary antibodies. After washing, the ligation and amplification steps were performed. Anti-mouse Alexa Fluor 488 was used as secondary antibody. Nascent DNA was detected as described above. Samples were mounted in ProLong™ Diamond Antifade Mountant.

### Confocal and widefield microscopy

Confocal images depicting MCF10A cells were acquired with a Leica TCS SP5 confocal laser-scanning microscope, using an HCX PL APO 100x/1.40/0.70 oil immersion objective lens (Leica Microsystems, Mannheim, Germany). Excitation was provided with a white laser, enabling the best condition as function of the utilized dye, namely: 488 nm (for Alexa Fluor 488), 532 nm (for Atto 532), 554 nm (for Quasar 579), 594 nm (for Alexa Fluor 594), and 647 nm (for Alexa Fluor 647 and Atto 647N), with relevant emission detection bands 495 – 525 nm (for Alexa Fluor 488), 540 – 620 nm (for Atto 532), 560 – 625 nm (for Quasar 570), 655 – 725 nm (for Alexa Fluor 647 and Atto 647N). Signals in different channels were detected with photomultipliers (PMT) or Hybrid Detectors (HyD). Typical pixel size is 40 nm. For images showing U2-OS cells a Zeiss LSM880 was used equipped with a Plan-Apochromat 40x/1.4 Oil DIC M27 objective and 405 nm (UV- photodiode), 488 nm (Argon ion), 561 nm (DPSS) and 633 nm (HeNe) laser lines. 12 bit images were recorded at a resolution of 1024×1024 pixels, with 2x digital zoom and a scanning speed of 7. Live cell imaging of MCF10A was performed with a NIKON A1R confocal microscope equipped with a stage top incubator and a hardware autofocus. Images were acquired in 1024 x 1024 pixel format, with a pixel size of about 85 nm, without averaging. Images were acquired every 5 min or 10 min over 10-12 hours. Excitation was provided with a CW Diode Laser (Coherent, Cube 488-50) at 488 nm for eGFP (DEK-eGFP) and a CW Diode Pumped Solid State Laser (DPSS, Melles Griot, 85YCA) at 561 nm for RFP (pCellCycleChromobody®-TagRFP). Widefield images of MCF10A cells were acquired on an AxioObserver or a CellObserver HS microscope (Zeiss) equipped with a PlanApochromat 40X/1.40 oil objective, a HXP 120 mercury arc lamp and an Axiocam MRm CCD camera (1300 × 1030 pixels). Fluorescence signals were detected with 493/517 nm (GFP and AlexaFluor 488), 590/612 nm (AlexaFluor 568) and 640/690 nm (AlexaFluor 647) filter sets. DEK bodies in U2-OS cells stably expressing eGFP-DEK or mutants thereof were imaged on a Celldiscoverer 7 widefield imaging platform (Zeiss) equipped with a Plan-Apochromat 20x/0.7 air objective and an Axiocam 512 camera. Binning was set to 2,2 and the exposure time to 200 ms. Cells were plated in 96-well imaging plates (Ibidi), 10 positions were recorded per well and images acquired every 12 min for a total duration of 22 h.

### STED microscopy

STED measurements were performed on a Leica TCS SP5 gated-STED microscope, using an HCX PL APO 100×100/1.40/0.70 oil immersion objective lens (Leica Microsystems, Mannheim, Germany). Emission depletion was performed with 592 nm CW STED laser. The 592nm STED laser power was 50% of 350 mW maximum power (about 175 mW). Excitation was provided with a white laser at 488 nm wavelength for Alexa Fluor 488 and with emission detection band 495 – 525 nm, with 1.50 ns - 9.50 ns time gating using a hybrid detector (Leica Microsystems). Typical pixel size is 40 nm, with the pixel dwell time of 2 μs and 48 lines averaging.

### SMLM Imaging

For the super-resolution SMLM imaging of DEK, MCF10A cells were washed with pre-warmed PBS 3x, and fixed with formaldehyde (3,7%, methanol free). After subsequent blocking with blocking buffer (0,1% Triton X-100 and 3% w/v of bovine serum albumin (BSA)) for 1h at room temperature, samples were incubated with the primary antibody (overnight at +4°C). F(ab’)2-Goat anti-Mouse IgG (H+L) Cross-Adsorbed Secondary Antibody, Alexa Fluor 647 was used as secondary antibody (ThermoFisher Scientific, A21237, 1:250, 45min in RT). Samples were post-fixed in PFA 2% for 5 min and stored in PBS at +4°C until applying GLOX buffer containing a glucose oxidase-based oxygen scavenging system for imaging (Bates et al., 2007). A commercial N-STORM & N-SIM TIRF Eclipse Ti2 microscope equipped with an oil immersion objective (CFI SR HP Apochromat TIRF 100XC Oil, NA 1.49) was used to acquire 20000 frames at a frame rate of 30 ms using oblique incidence excitation and sCMOS camera (ORCA-Flash4.0, Hamamatsu Photonics K.K.). Alexa Fluor 647 labeled samples were switched into dark state with the 647 nm laser and reactivated with the 405 nm laser. Imaging cycles consisted of 1 activation frame followed by 3 read-out frames. Acquisition was performed using the continuous STORM filter and hardware autofocus. Image reconstruction was performed with the NIS-Elements and N-STORM Analysis imaging software.

### Fluorescence Recovery After Photobleaching

FRAP measurements were performed on a Leica TCS SP5 confocal laser scanning microscope. The experiments were conducted under a controlled environment in a dedicated live-cell imaging chamber (5 % of CO_2_, 37°C, 90 % humidity). Briefly, regions of interest (ROI) of 2 μm x 2 μm were selected and photobleached with 100% laser power (55 μW @488nm). Subsequently, fluorescence recovery was monitored for about 3 min, all traces were min/max normalized, averaged and the standard error was calculated. The parameters used for the FRAP measurement are listed in Table 1.

**Table 1.** Parameters used in FRAP experiments and data acquisition.

|  | DEK-eGFP |
| --- | --- |
| FRAP image acquisition | 512 x 512 pixels |
|  | 40 nm pixel size |
|  | Excitation wavelength: 488 nm 10% |
|  | Emission: 500 nm - 580 nm |
|  | 1400 Hz scan speed |
| Pre-photobleaching | 2 frames (time/frame 0,37 s) |
| Photobleaching protocol | 5 frames (time/frame 0,37 s);<br>2 x 2 µm square ROI; 100% 488 nm laser |
| FRAP imaging | 90 frames (time/frame 2s); 10% 488 laser |

### Image analysis

Image analysis was performed using ImageJ (http://imagej.nih.gov/ij/) and Leica LAS X F (Leica Microsystems GmbH). The number of DEK bodies was obtained from Z-stack Maximum projections on which was performed Find Maxima function (with the prominence about 80-90 depending on the level of immunofluorescence staining).

For finding the number of DEK bodies colocalizing with PCNA DNA replication foci, the object-based analysis was performed by using an ImageJ plugin JaCoP (Bolte and Cordelieres, 2006). For the analysis of FRAP data, first the photobleached region within cell nucleus was cropped. After this, the mean intensity of the ROI along the time experiment was plotted, and the curves were combined and normalized to the minimum and maximum of the combined curve. Finally, each of the curves was fit. The tau parameter was obtained from the average of all fits. This procedure was performed for the case of DEK within DEK bodies and for DEK within the other chromatin regions.

### Preparation of cell extracts, immunoprecipitation, and immunoblot analysis

Whole cell lysates were prepared *via* scraping cells in PBS containing protease inhibitors (Complete Protease inhibitor cocktail, Roche). Cells were pelleted and resuspended in hot SDS lysis buffer (0.5 % SDS, 50 mM Tris pH 7.5), incubated for 10 min at 95°C, centrifuged, resulting samples were frozen in liquid nitrogen and stored at -80°C.

For preparation of cell lysates for immunoprecipitation, cells were harvested in hypotonic buffer (20 mM HEPES pH 7.4, 20 mM NaCl, 5 mM MgCl_2_, 10 mM NaF, 1 mM Na-vanadate, 1 mM PMSF, 1x Complete Protease Inhibitor Cocktail (Roche)) supplemented with 10 mM N-Ethylmaleimide (NEM). After centrifugation (5 min, 500 x g, 4 °C), the resulting cell pellet was resuspended in hypotonic buffer to 1 x 10^6^ cells per 100 μl. For extraction of chromatin-bound DEK, 450 mM NaCl and 0.5% NP-40 was added, incubated for 15 min on ice and harvested at 10,000 x g for 10 min at 4 °C.

For preparation of cell lysates for nickel affinity chromatography (6xHis-SUMO pulldown), a previously published protocol was carried out (Tatham et al., 2009). HeLa cells stably expressing 6xHis-SUMO-1 (accession number P62165), 6xHis-SUMO-2, (accession number CAG46970) or 6xHis-SUMO-3 (accession number CAA67897) as well as HeLa Rachel cells as negative control were grown in 10 cm cell culture dishes to 80% confluency. Cells were harvested *via* scraping in 5 ml PBS containing 10 mM NEM. Crude cell extracts for visualization of SUMO conjugates were prepared with 1 ml of cell suspension. Crude samples were centrifuged (2 min, 1000 x g, RT), the supernatant was removed, and the pellet was resuspended in 200 μl 2x Laemmli sample buffer (200 mM imidazole, 5% SDS, 150 mM Tris-HCL pH 6.7, 30% glycerol, 0.0025% bromophenol blue). Samples were boiled for 5 min at 95°C, then mechanically disrupted by passing through a 20G needle 20 times using a 1-ml syringe and supplemented with 750 mM β-mercaptoethanol. The remaining 4 ml cell suspension was centrifuged (5 min, 3000 x g, RT) and the pellet was dissolved in 5 ml of cell lysis buffer (6 M Guanidinium-HCl, 10 mM Tris, 100 mM sodium phosphate buffer pH 8.0, 0.1% Triton X-100, 10 mM NEM). Samples were supplemented with 5 mM β-mercaptoethanol and 5 mM imidazole and sonicated on ice for 40 s at medium power. Cell debris was removed by centrifugation (15 min, 3000 g, RT) and the supernatant was stored at -20 °C.

For immunoprecipitation all buffers were supplemented with 10 mM NEM. 6 μl of polyclonal rabbit α-DEK K-877 (Kappes et al., 2008) was used for 100 μl cell lysate. After 30 min incubation on ice, 100 μl of a 25 % Protein A/G Plus-Agarose (SantaCruz) was added and rotated for 1 h at 4 °C. After washing three times with extraction buffer (20 mM HEPES pH 7.4, 300 mM sucrose, 450 mM NaCl, 0.5 mM MgCl_2_), immune complexes were eluted from the beads with 2 % SDS and 5 % β-mercaptoethanol for 1 h at 37 °C. IP samples were concentrated using methanol-chloroform protein precipitation (Wessel and Flügge, 1984), solubilized in 1x SDS sample buffer and boiled for 10 min at 99 °C. Samples were separated by SDS-PAGE, blotted to nitrocellulose and analyzed by immunoblotting.

For nickel affinity chromatography for purification of 6xHis-SUMO proteins all buffers were supplemented with 10 mM NEM. The prepared cell lysates were subjected to chromatography using HisPur Ni-NTA Resin (Thermo Fisher Scientific). The lysate was added to 100 μl packed volume of beads per sample. Bead slurry was pre-washed three times with ten bead volumes of cell lysis buffer (6 M Guanidinium-HCl, 10 mM Tris, 100 mM sodium phosphate buffer pH 8.0, 0.1% Triton X-100). After tumbling the mixture overnight at 4°C, the bead suspension was centrifuged (2 min, 750 g, RT) and the supernatant carefully removed. Beads were washed once with 4 ml of cell lysis buffer containing 5 mM β-mercaptoethanol and once with 4 ml of 8.0 pH wash buffer (8 M Urea, 10 mM Tris, 100 mM sodium phosphate buffer pH 8.0, 0.1% Triton X-100, 5 mM β- mercaptoethanol). Subsequently, the beads were washed three times with 4 ml of 6.3 pH wash buffer (8 M Urea, 10 mM Tris, 100 mM sodium phosphate buffer pH 6.3, 0.1% Triton X-100, 5 mM β-mercaptoethanol). Finally, the beads were resuspended in 1.5 ml of 6.3 pH wash buffer and then transferred to 1.5 ml centrifuge tubes. The wash buffer was completely removed after centrifugation (1 min, 6000 x g, RT) and elution of 6xHis-SUMO proteins was performed with 100 μl of elution buffer (200 mM imidazole, 5% SDS, 150 mM Tris-HCL pH 6.7, 30% glycerol, 720 mM β-mercaptoethanol, 0.0025% bromophenol blue) per sample. The beads were boiled for 2 min at 95°C and then incubated for 20 min at RT. The eluted fraction and the crude cell extracts were separated by SDS-PAGE, blotted to nitrocellulose and analyzed by immunoblotting.

For analysis of whole cell lysates 40 μg of protein were loaded onto a 12 % SDS gel and separated by electrophoresis. Proteins were transferred onto a nitrocellulose membrane using semi-dry Western blotting.

For analysis of eluted 6xHis-SUMO proteins 100 μl of eluted fraction and input/crude samples were loaded onto a 10 % SDS gel and separated by electrophoresis. Proteins were transferred onto a nitrocellulose membrane using wet Western blotting.

DEK-IP eluates were separated on a 12.5 % SDS gel. Membranes were blocked in 5 % milk/TBS-T and incubated with primary antibodies overnight at 4 °C under mild agitation (anti-DEK K-877). Membranes were washed and incubated with HRP-conjugated secondary antibodies for 1 h at RT (Goat-anti-Rabbit-HRP, Goat-anti-Mouse-HRP, both Dako Denmark). Marker lanes were incubated with Streptavidin-biotinylated-HRP (GE Healthcare). Proteins were visualized using chemiluminescent detection with ECL Prime Western Blotting Detection Reagents (Amersham RPN2232).

### Site-directed mutagenesis

Nucleobase mutations of the DEK sequence were introduced using an improved QuickChange site-directed mutagenesis protocol (Zheng et al., 2004). The pEGFP-N1- hDEK plasmid (DEK WT sequence inserted into eGFP reporter plasmid “peGFP-N1”; Addgene 6085-1) was used as a template for mutagenesis PCR. The potential SUMOylation site 1 (bases 181 – 186) and the SUMO consensus motif (bases 778 – 789) were mutated using overlapping primers containing the desired base change. For the SUMOylation site 1 two codons were mutated (lysines 61 and 62 to alanine) in two rounds of mutagenesis. For the SUMO consensus motif one round of mutagenesis was performed (lysine 261 to alanine). For SUMO consensus motif mutation, the plasmid DNA was denatured prior to the PCR using 0.2 M NaOH to reduce the amount of colonies containing unmutated wildtype-plasmid after transformation. Primer sequences are listed in **Supplementary Table S4**.

### Expression and purification of recombinant GST-tagged proteins from *E*. *coli*

The mutated DEK sequences (DEK SUMOmut 61-62, 144, 145 or 261) were inserted into a GST expression plasmid (pGEX 4T-1), transformed into *E*.*coli* BL21(DE3) (Thermo Fisher Scientific) and purified as described before (Ganz et al., 2019). Briefly, protein expression was induced with 0.5 mM IPTG (Sigma-Aldrich) for 1.5 h. Bacteria were harvested *via* centrifugation at 4600 g for 15 min at 4°C and resuspended in resuspension buffer (20 mM Tris-HCl (PH 8.0), 1 M NaCl, 0.5 mM EDTA, 1 mM DTT) and frozen in liquid nitrogen. After thawing, bacteria were sonicated on ice and 0.5 % NP-40 (Sigma-Aldrich) was added and centrifuged at 18000g for 30 min at 4°C. The supernatant was added to 200 μl Glutathione-Sepharose 4B-beads (GE Healthcare) and incubated for 2 h at 4°C. After centrifugation, beads were washed with wash buffer with decreasing NaCl concentrations (500 mM, 300 mM, 20 mM) and GST-tagged DEK was eluted with 200 μl elution buffer (200 mM Tris-HCl (pH 8.0), 200 mM NaCl, 40 mM reduced glutathione (Sigma-Aldrich), 10 % glycerol) for 1 h at 4°C. The supernatant was frozen in liquid nitrogen and stored at -80°C.

### In vitro SUMOylation

For in vitro SUMOylation, recombinant SUMO E1, SUMO E2 (UBC9), SUMO-1 and SUMO-2 were purchased from Enzo Life Sciences (Lausen, Switzerland). Reactions were performed for 2 hours at 30°C in *in vitro* buffer (20 mM Tris-HCl (pH 7.6), 50 mM NaCl, 10 mM MgCl2, 0.1 mM dithiothreitol (DTT), 1x protease inhibitor cocktail (cOmplete™, Mini, EDTA-free Protease Inhibitor Cocktail, Roche) with or without 4 mM ATP) as described before (Aichem et al., 2019). Reaction was terminated by addition of 5x SDS sample buffer with 4 % β-mercaptoethanol and boiled for 5 min at 95°C. 20 μl reactions were mixed containing 0.71 μg recombinant protein (GST, GST-DEK, GST-DEK SUMOmut 61-62, GST-DEK SUMOmut 261 or GST-DEK Sumo mut 61,62,144, 145, 261), 450 ng SUMO E1, 1 μg SUMO E2, 3 μg SUMO-1 or SUMO2.

## Acknowledgements

The authors thank Mario Faretta (European Institute of Oncology, Milan, Italy) for MCF10A cells; Elena Cerutti (Istituto Italiano di Tecnologia -IIT, Genova, Italy) for help with RNA-FISH; Annette Aichem (Biotechnology Institute Thurgau, Tägerwilen Switzerland) for help with *in vitro* SUMOylation and Christian Tischer (EMBL Heidelberg, Germany) for help with image data analysis; the core facilities FlowKon and Bioimaging Center at the University of Konstanz (Germany) for assistance in microscopy and flow cytometry experiments and the Advanced Light Microscopy Facility (ALMF) of EMBL for hosting and supporting the primary siRNA screen; Michele Oneto (IIT), Marco Scotto (IIT) Luca Lanzanò (University of Catania, Italy) and Martin Stöckl (University of Konstanz) for fruitful discussions.

## Data availability

R code used for DEK body analysis is provided at https://github.com/CP-Vogel/R-DEK-body-script.git.

## SUPPLEMENTAL INFORMATION

**Supplementary Figure S1:**
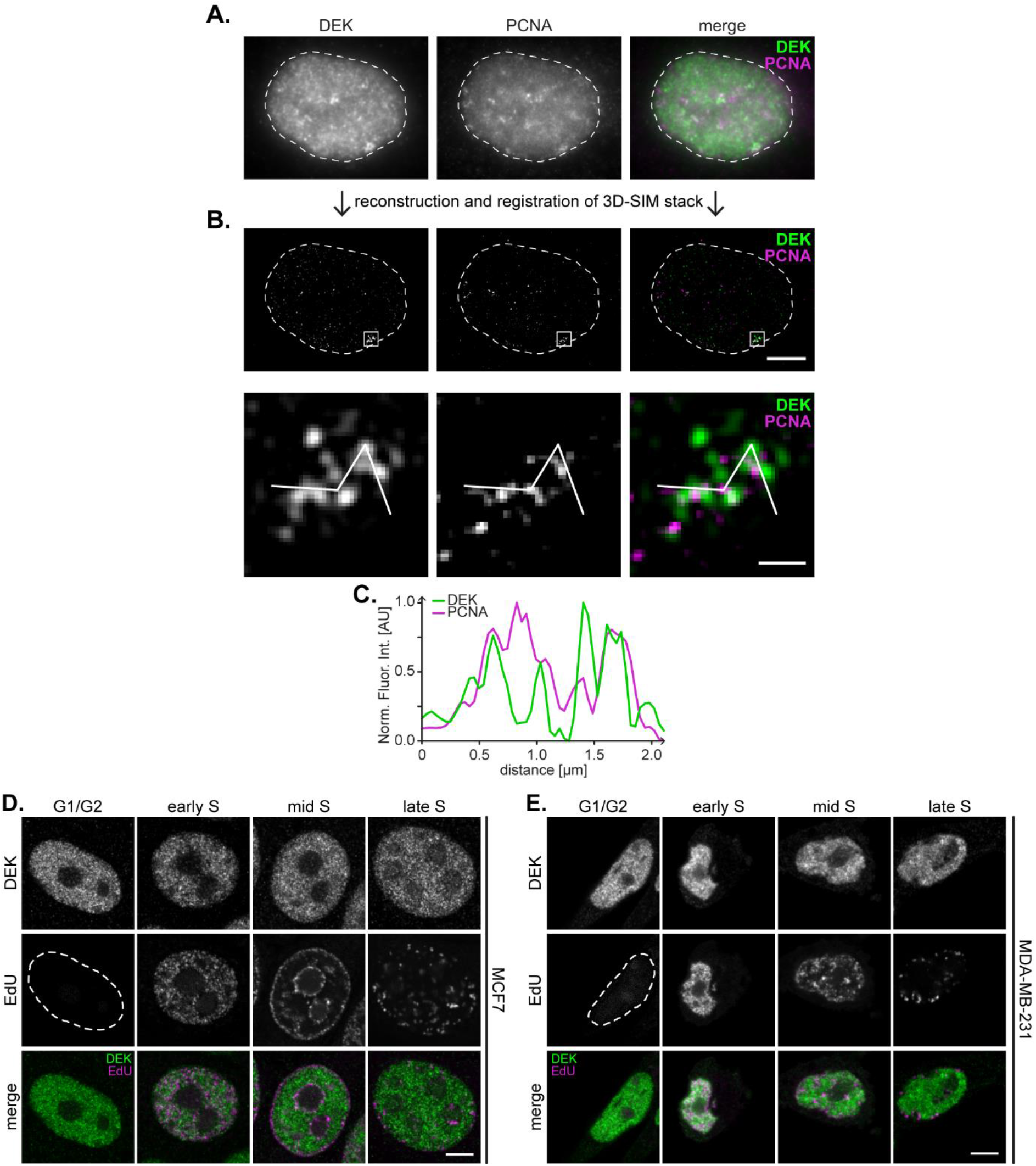
Endogenous DEK forms DEK foci in primary BJ5-ta cells, yet not in malignant MCF7 and MDA-MB-231 cells. **A.** Immunofluorescence analysis of DEK (green) and PCNA (magenta) distribution in a BJ5-TA fibroblast nucleus by super-resolution microscopy (3D-SIM) *via* indirect immunofluorescence. Pseudo-widefield representation of one slice. **B.** Image of a single Z-slice from a middle section of the super-resolved image stack after reconstruction and registration. Scale bar: 5 μm. The magnified inset shows one DEK- positive large replication focus. Scale bar: 0.5 μm. **C.** Fluorescence intensity profiles of DEK and PCNA. Interpolated intensity profiles were calculated along the line indicated in (B) and normalized to the min/max values. **D, E** Occurrence of DEK bodies is reduced in (D) MCF7 and (E) MDA-MB-231 cells. Immunofluorescence analysis of DEK and EdU distribution pattern in late S-phase cells was performed as described in Fig. 1. Scale bar: 5 μm.

**Supplementary Figure S2:**
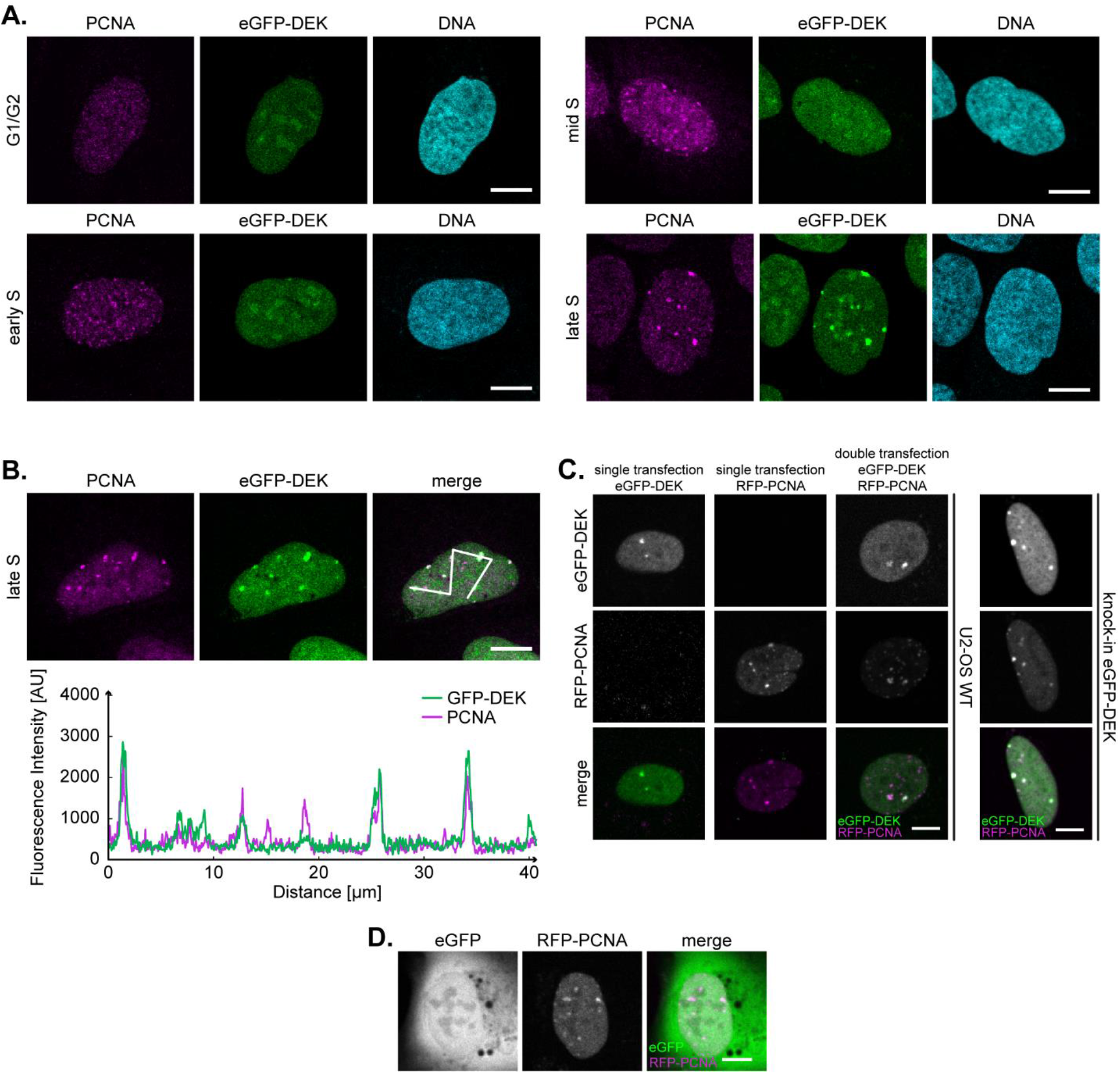
Colocalization of eGFP-DEK and PCNA during late S- phase in U2-OS cells carrying a TALEN-mediated eGFP-DEK genomic knock in. **A.** Representative confocal images of U2-OS KI eGFP-DEK cells in different cell cycle phases labeled with PCNA-specific antibodies. Similar focal accumulations of eGFP-DEK (green) and PCNA (magenta) can be observed in late S-phase. DNA was counterstained with Hoechst33342 (cyan). Scale bar: 10 μm. **B.** Colocalization analysis of PCNA and DEK bodies in U2-OS KI eGFP-DEK cells. Scale bar: 10 μm. Representative images are shown, and intensity profile was analyzed along the white line displayed in the right panel. **C.** DEK body formation is independent of the DEK expression level. U2-OS wild type cells were transfected with plasmids either coding for pEGFP-N1-hDEK or pENmRFP- PCNAL2 or double transfected with both plasmids and incubated for 24 h for maximal protein expression. U2-OS KI eGFP-DEK cells were transfected with pEN-mRFP- PCNAL2 only. Cells were monitored for 24 h (37°C, 5% CO2). Concomitant with DEK foci formation, relocation of RFP-PCNA (magenta) can be detected both in transiently transfected cells and in the stable cell line. Single transfected cells (upper two panels) serve as control cells. Scale bar: 10 μm. **D.** DEK foci formation is not an eGFP artifact. U2-OS cells were double transfected with pEGFP-C1 and pENmRFP-PCNAL2 and incubated for 24 h. During RFP-PCNA body formation, an equally distributed eGFP signal can be visualized. Scale bar: 10 μm.

**Supplementary Figure S3:**
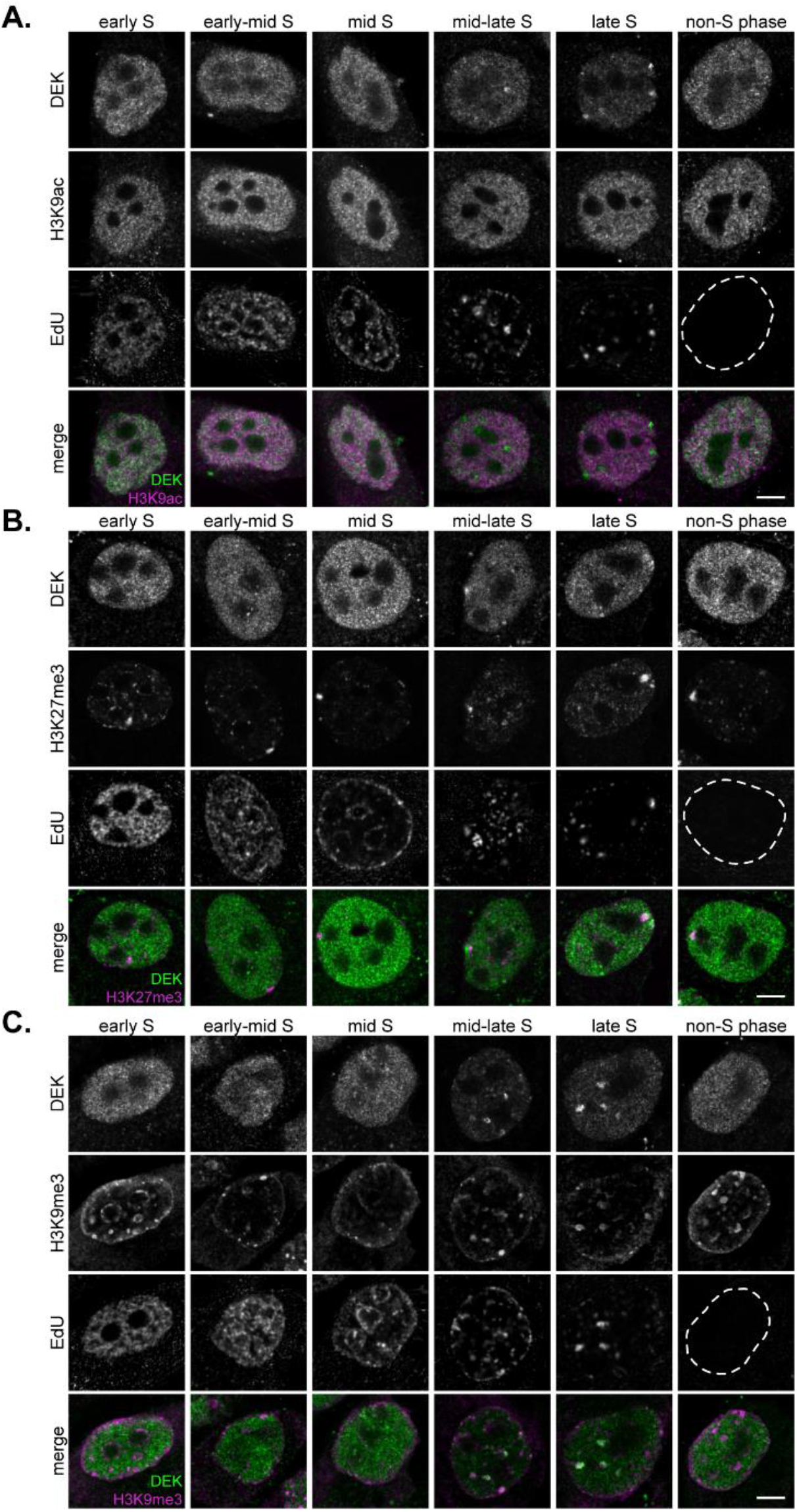
Localization analysis of DEK and histone marks in S- phase (data set corresponding to Fig. 3A). Confocal images of MCF10A cells labeled with antibodies specific for DEK and histone post-translational modifications H3K9ac (A), H3K27me3 (B), and H3K9me3 (C) during S-phase. Scale bars = 5 μm.

**Supplementary Figure S4:**
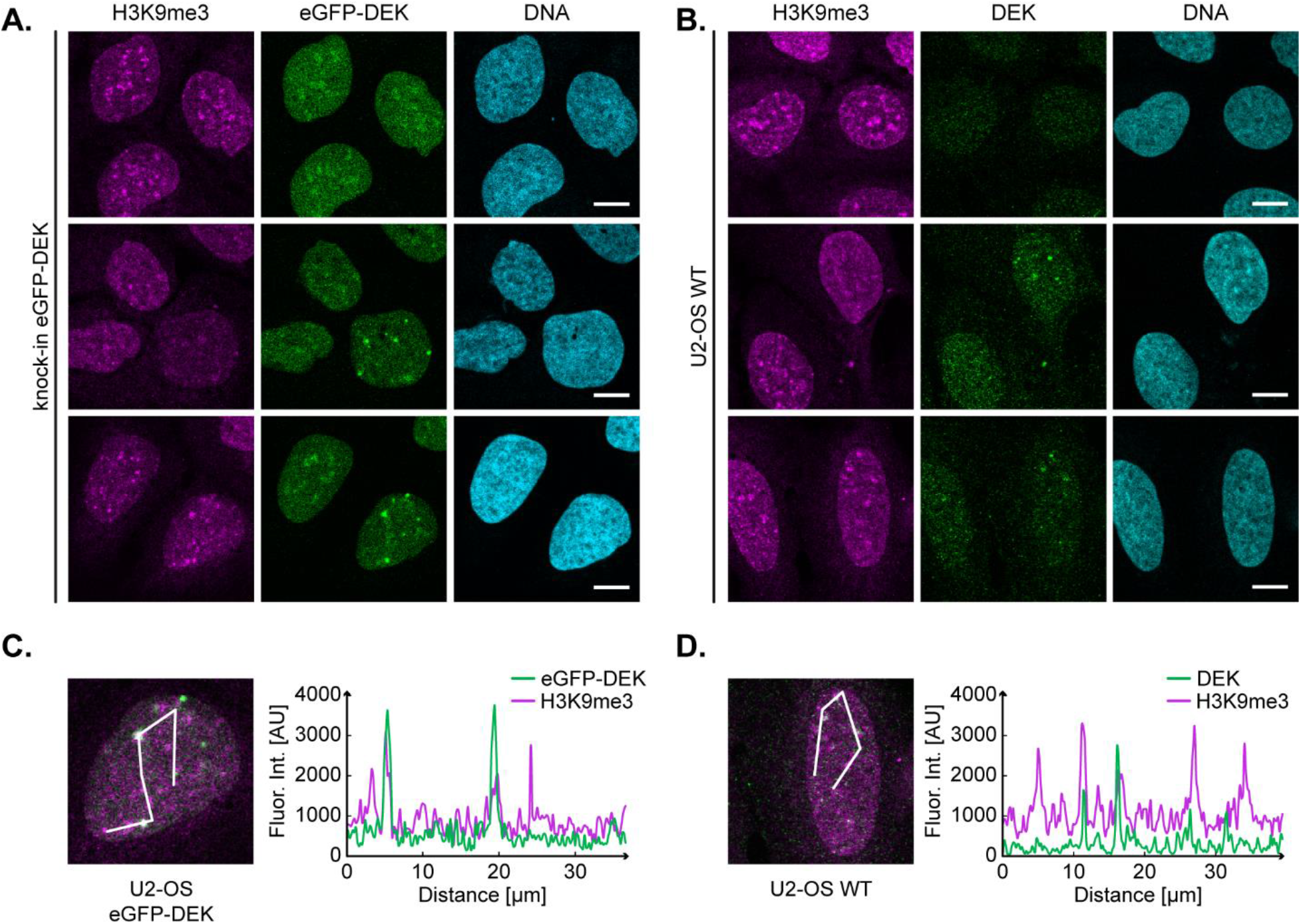
DEK bodies co-localize with heterochromatin in U2-OS cells. **A, B**. Representative confocal images of U2-OS KI eGFP-DEK labelled with H3K9me3 specific antibodies (magenta) (A) and of U2-OS WT cells (B) additionally labelled with antibodies specific for DEK (green). DNA was counterstained using Hochst33342 (cyan). Scale bar: 10 μm. **C, D**. Confocal images of single U2-OS KI eGFP-DEK and U2-OS WT cells from (A) and (B) with the respective fluorescence intensity profiles for the analysis of H3K9me3 (magenta) and DEK (green) colocalization. The white line is given as distance in the graph.

**Supplementary Figure S5:**
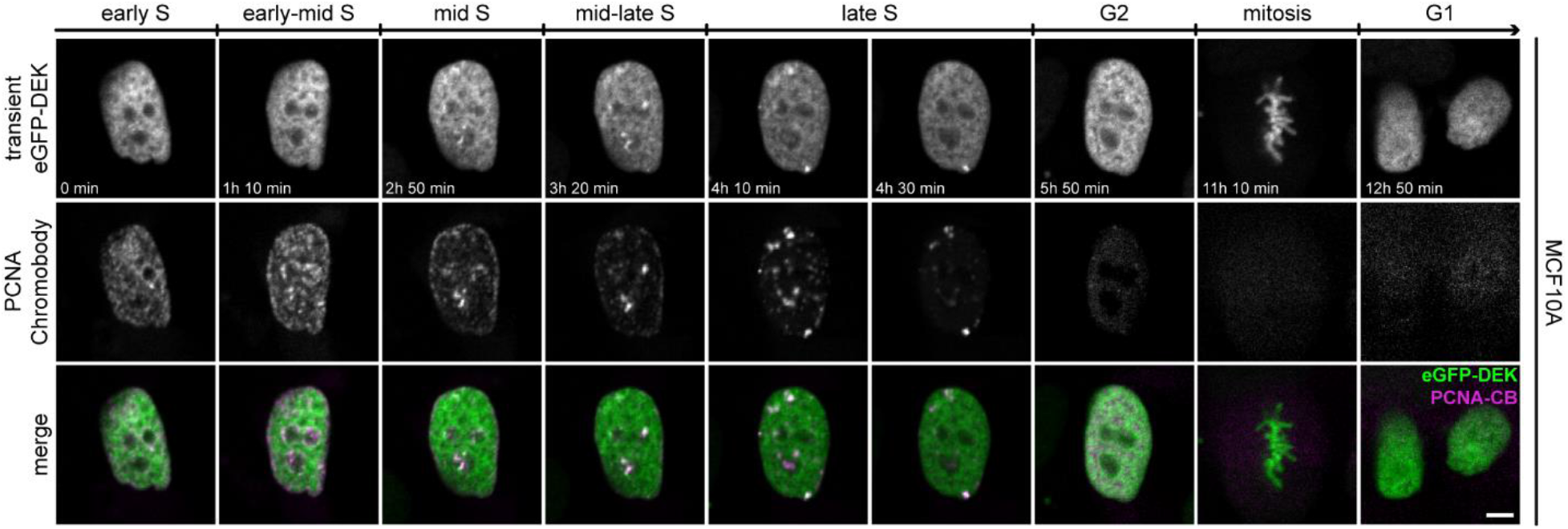
DEK bodies form during late S-phase of the cell cycle in transiently transfected MCF10A cells. Time-lapse fluorescence imaging of MCF10A cells co-transfected with plasmids encoding eGFP-DEK and mRFP-PCNA. The images were acquired at a scanning confocal microscope. Frames were captured every 10 min for 12h. DEK bodies can be traced for about 189 ± 38 min (13 measurements from 4 independent samples).

**Supplementary Figure S6:**
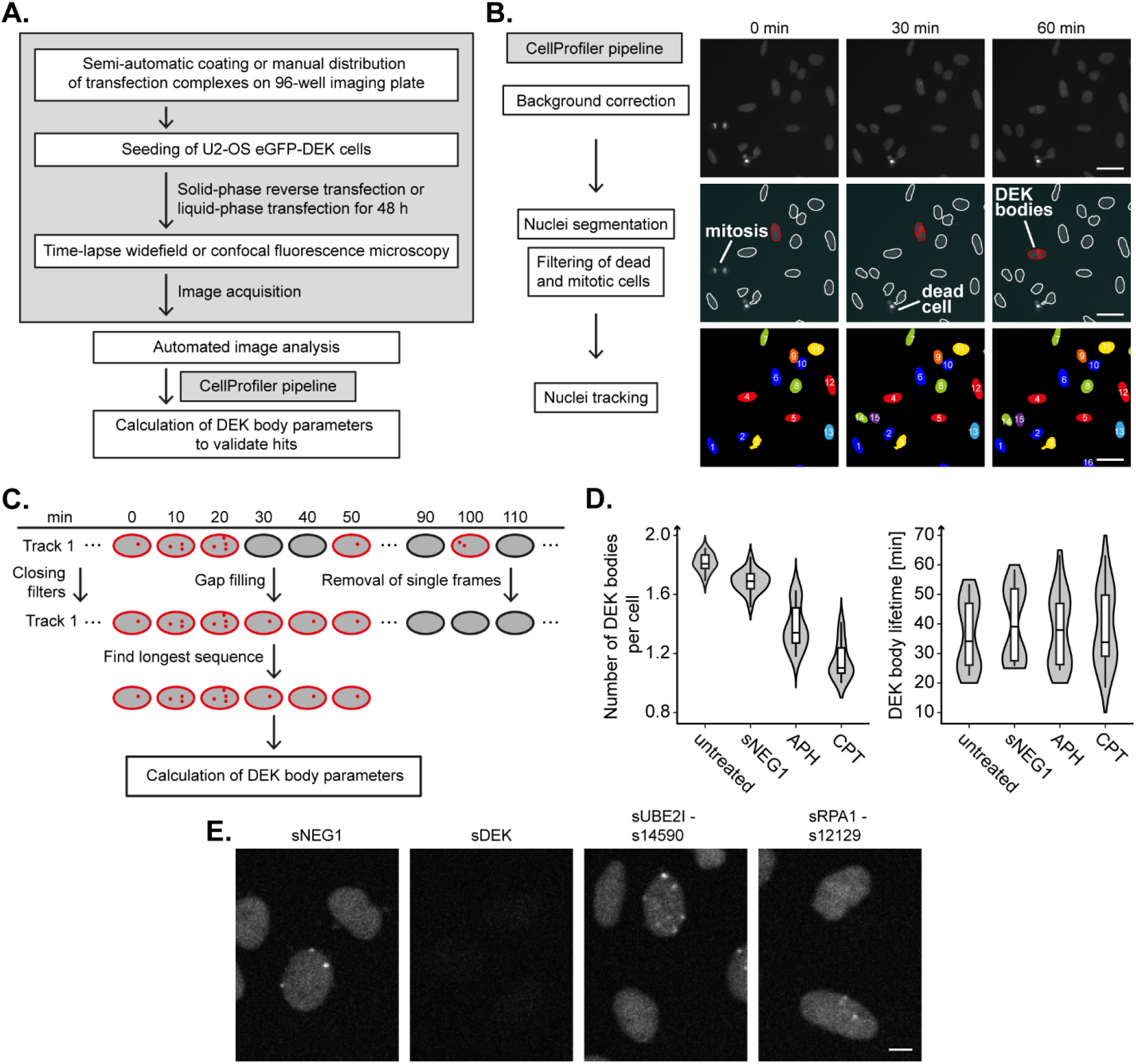
Design of a high-throughput siRNA screening approach for the identification of DEK body regulators. **A.** Schematic of the experimental protocols used for the siRNA screens. The pilot and primary screens relied on automated procedures and solid-phase reverse transfection of the siRNAs. Imaging was performed with a high-content widefield microscope. For the validation screen, cells and transfection reagents were dispensed manually. Imaging was performed on a spinning disk confocal microscope with an automated stage. Two pilot screens with slightly different imaging conditions were performed to optimize imaging parameters. The primary screens were performed in duplicate, and the validation screen in triplicate. **B.** Main steps of the image analysis pipeline. Example images of cells transfected with a negative control siRNA are shown. Nuclei with detected DEK bodies are outlined in red, nuclei that passed quality control in white. Nuclei that were rejected because of cells were dead or mitotic are not outlined. DEK bodies are encircled in red. Scale bar: 20 μm. **C.** Scheme of the gap-filling algorithm used after image analysis. The algorithm removed frames in which detection of DEK bodies within one track failed (false negatives), and sorted out frames in which foci were detected in one frame only (false positives). **D.** Quantification of DEK body parameters in image series from one of the pilot screens processed as in (A) and (C). Calculations were done using R. Violin plots display the mean number of DEK bodies per cell (left panel), and the length of the time windows in which DEK bodies appeared (DEK body lifetime, right panel) for the indicated treatments/siRNAs. In total, image data from 12 fields of view (4 positions per well, 3 wells per condition), each containing several tracks, were evaluated. Values from tracks within a field of view were averaged. The violin plot represents the density distribution of these averages, the black line indicates the median, the box the interquartile range (IQR) and the whiskers the IQR ± 1.5. **E.** Representative exemplary images of U2-OS KI eGFP-DEK cells treated with sNEG1, sDEK and the DEK body up-regulating siRNA sUBE2I and the down-regulating siRNA sRPA1. Images were taken at a spinning disk confocal microscope. Scale bar: 10 μm.

**Supplementary Figure S7:**
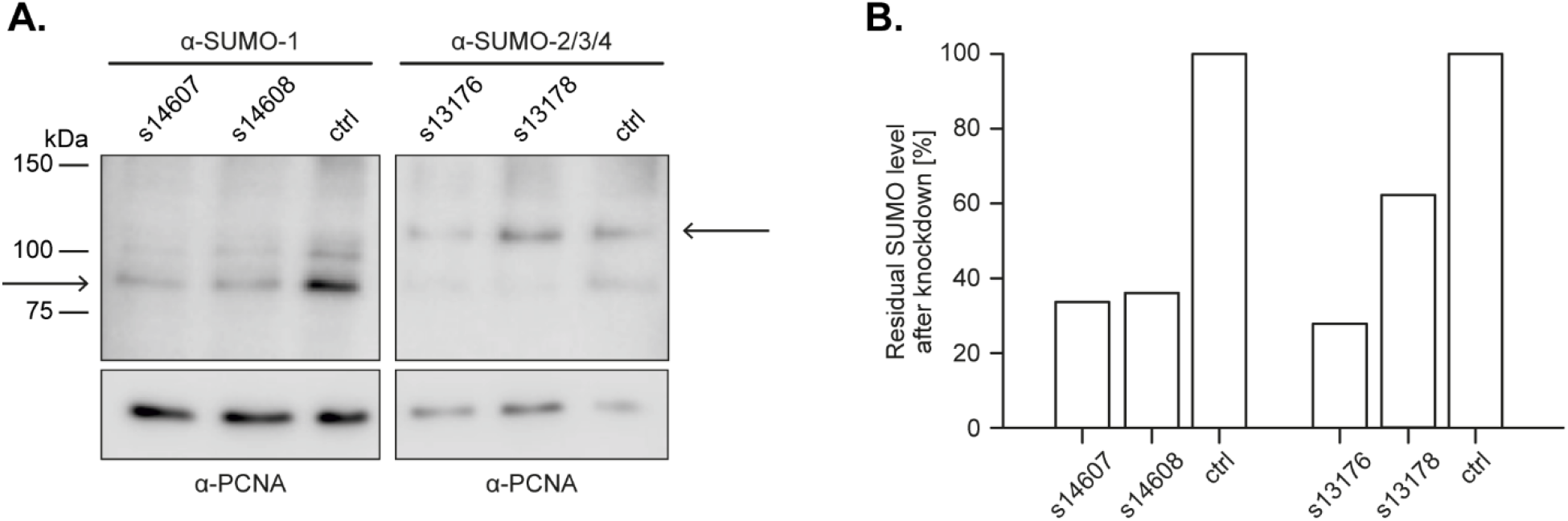
SUMO1/3 siRNAs are effective in down-regulating SUMO1/3 protein levels. **A.** Western blot analysis of SUMO1 and SUMO2/3/4 protein levels after siRNA-mediated knockdown of gene expression. Whole cell lysates from siRNA-transfected U2OS KI eGFP-DEK cells were subjected to Western blotting using SUMO1- and SUMO2/3- specific antibodies, respectively. As negative control the sNEG1 siRNA was used, PCNA served as loading control. Quantified SUMO signals are marked with an arrow. The experiment was performed once. **B.** Densitometric quantification of (A). SUMO-specific signals were normalized to PCNA and are plotted as percentage of their respective controls (PCNA).

**Supplementary Figure S8:**
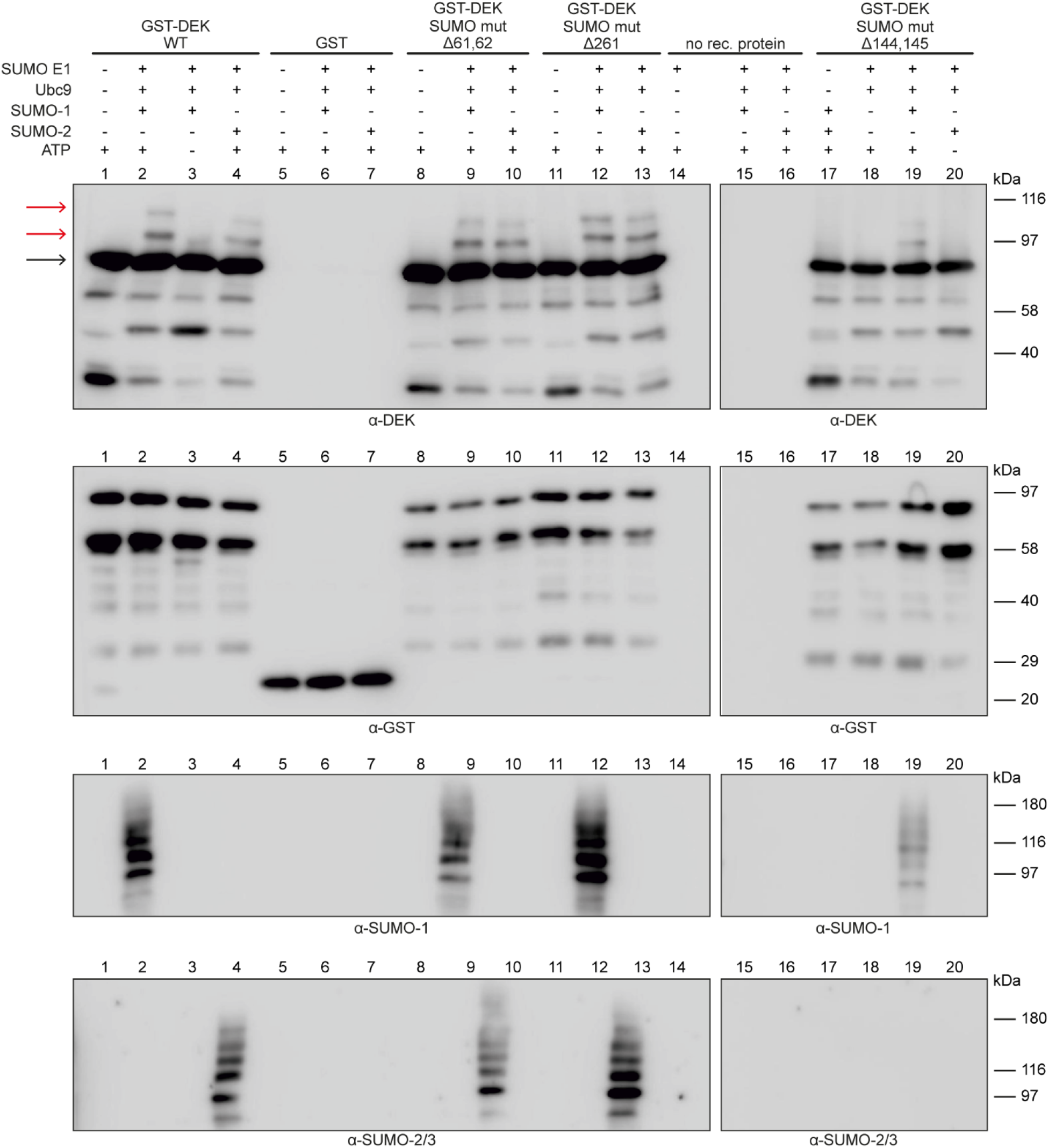
Mutation of individual DEK SUMOylation sites does not lead to reduced DEK SUMOylation in vitro. Samples containing recombinant GST-tagged DEK WT, mutant, or GST only proteins as indicated were subjected to *in vitro* SUMOylation assays as described in Fig. 8. Proteins were detected with DEK-, GST-, and SUMO-specific antibodies. The black arrow indicates GST-DEK signals, red arrows the mono and di-SUMOylated variants.

**Supplementary Table 1.**
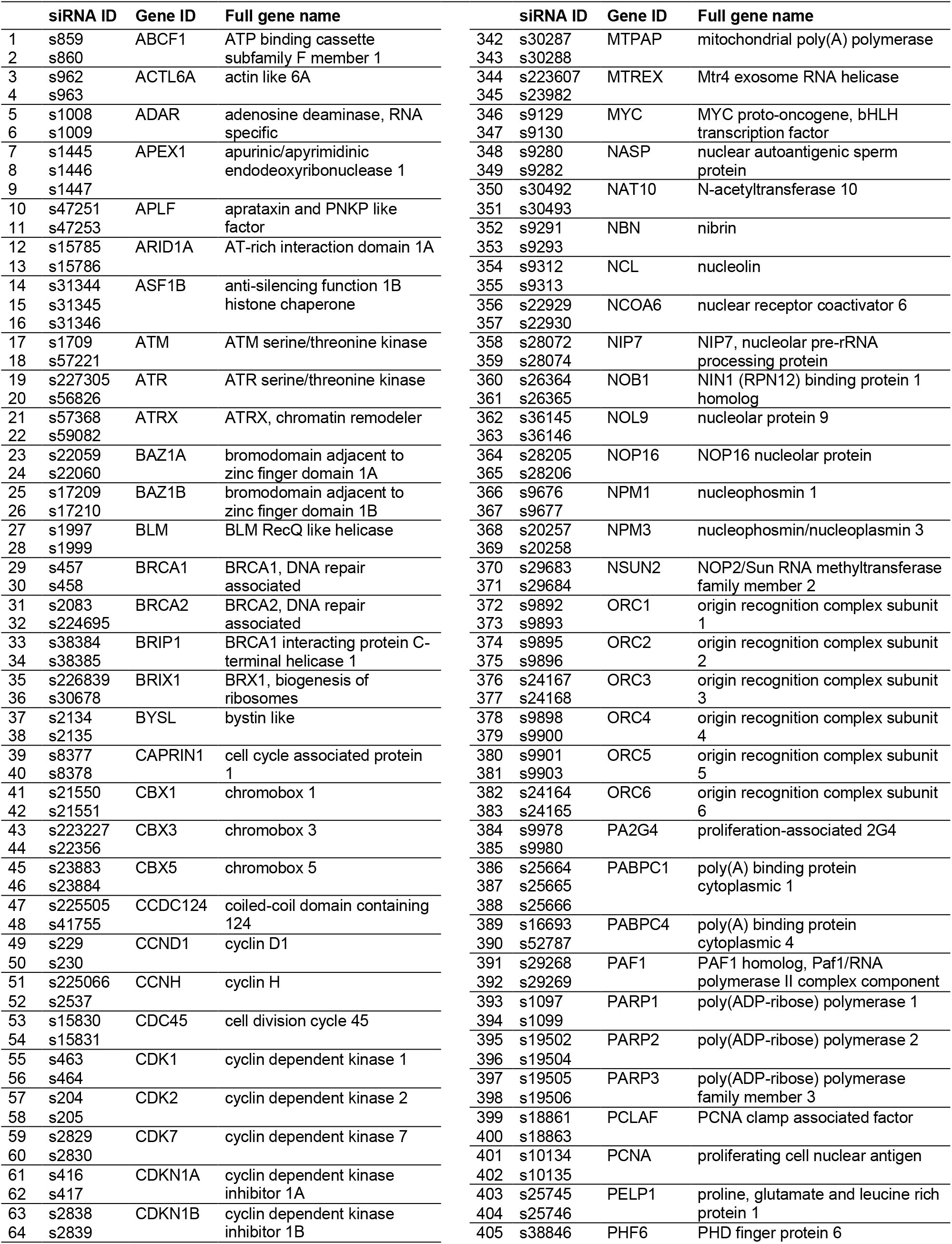
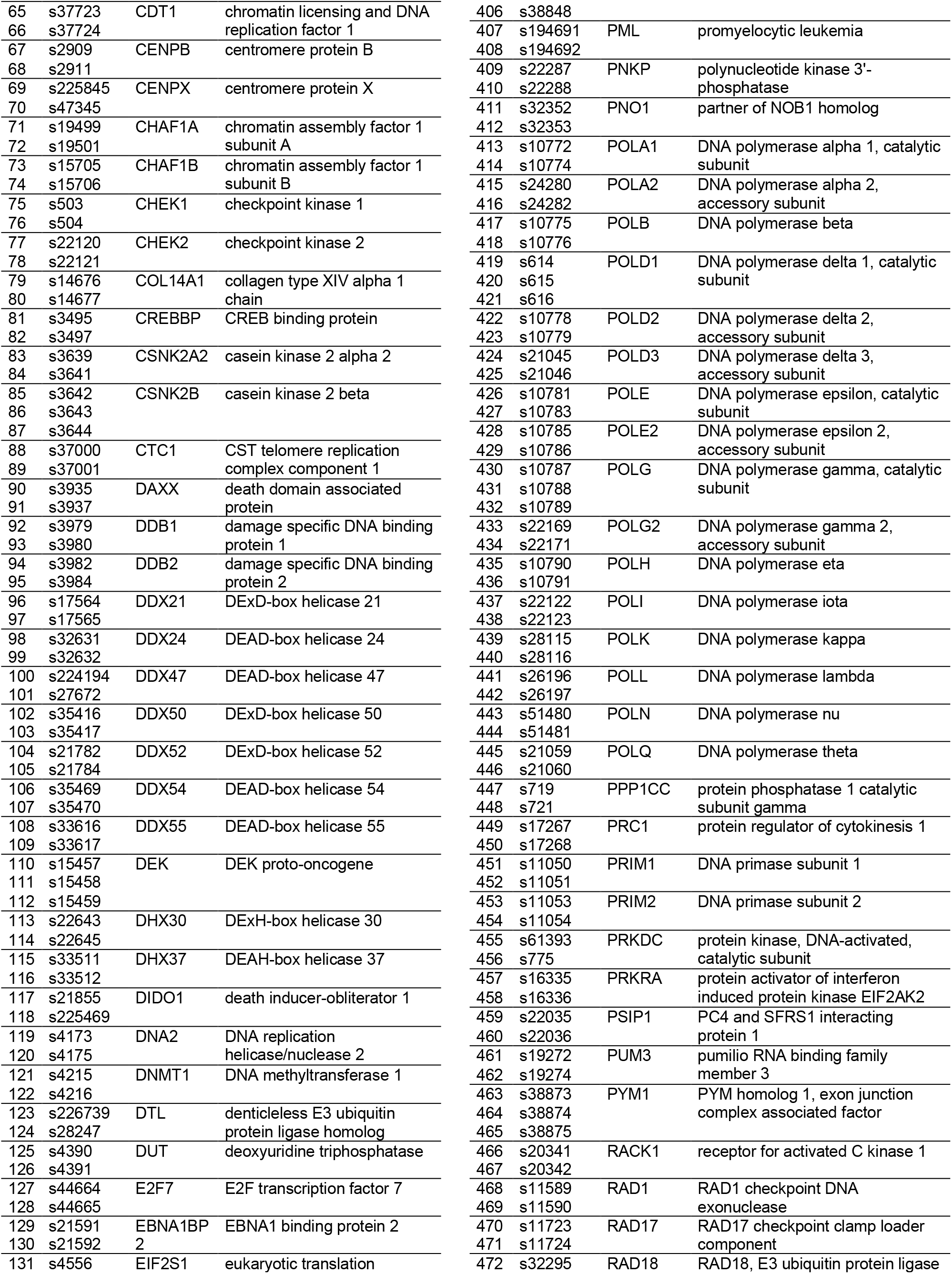
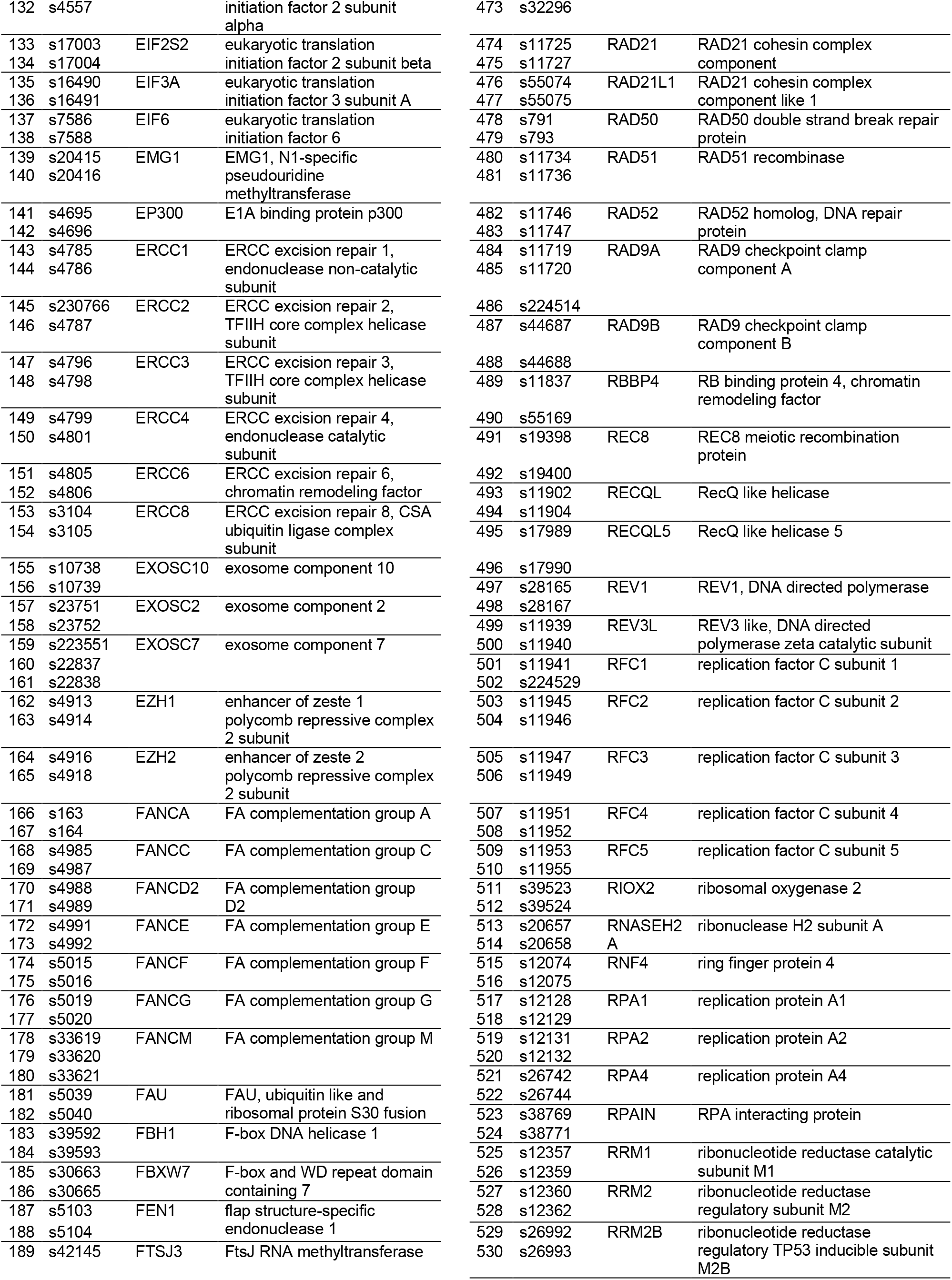
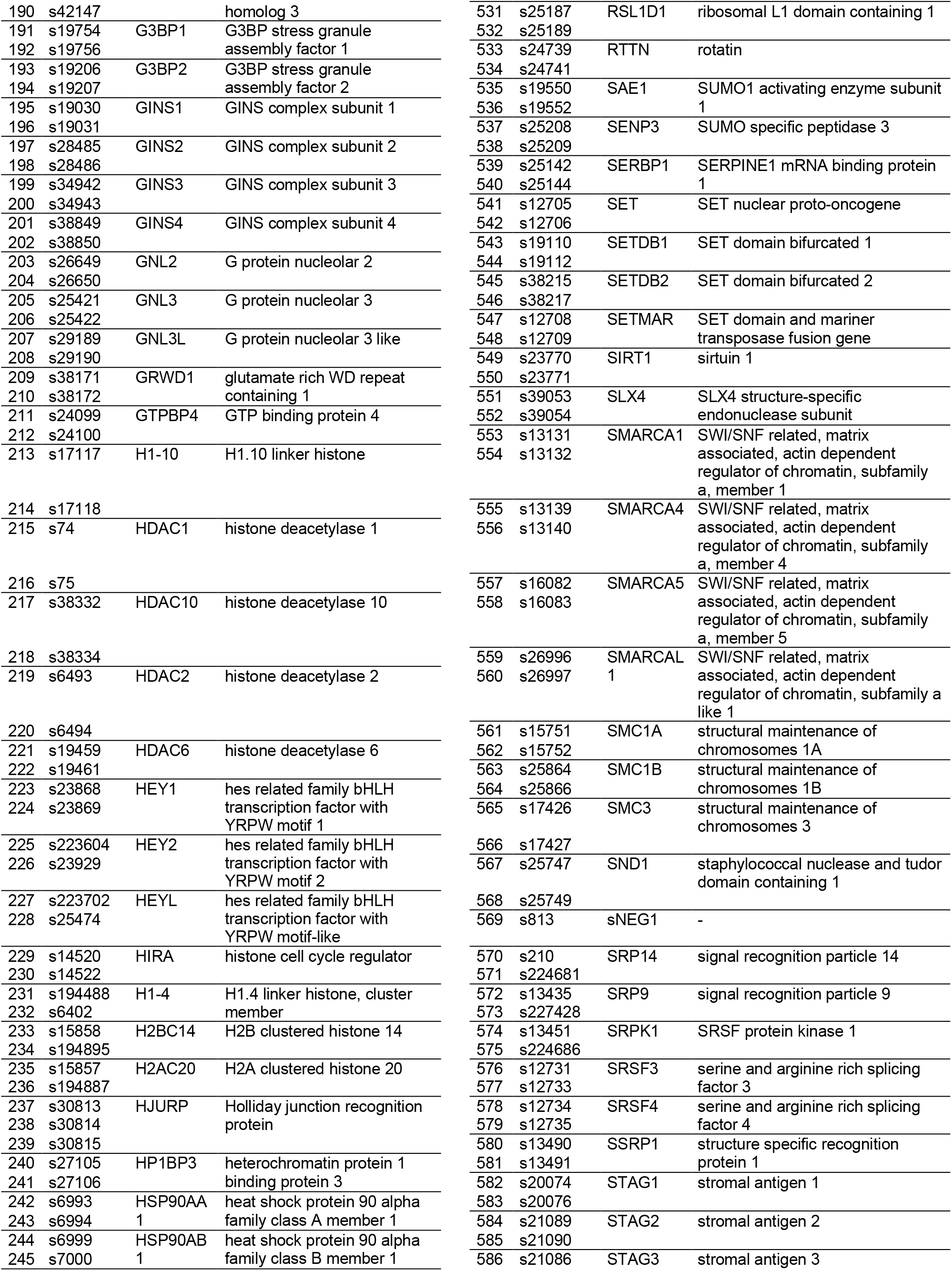
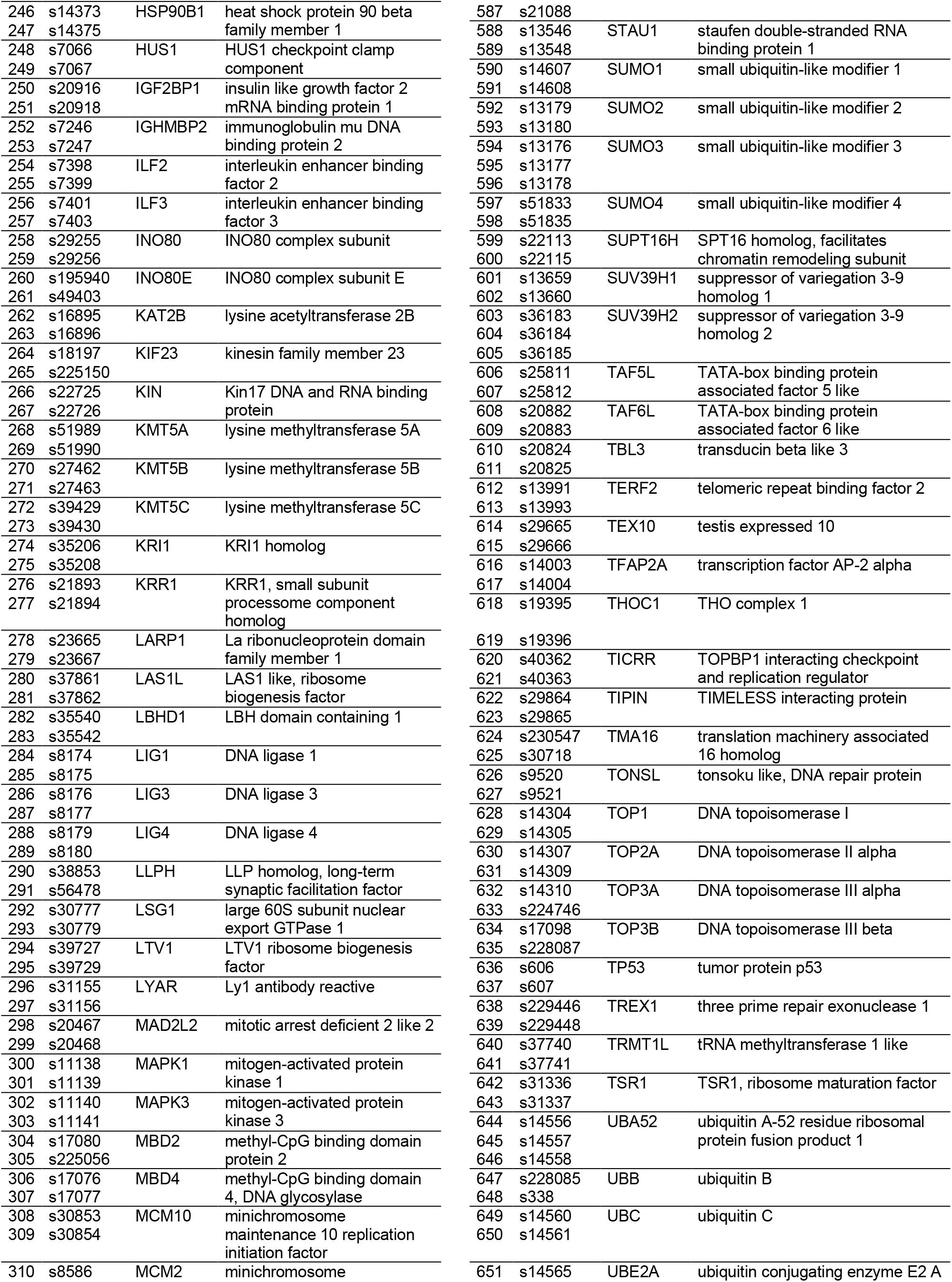
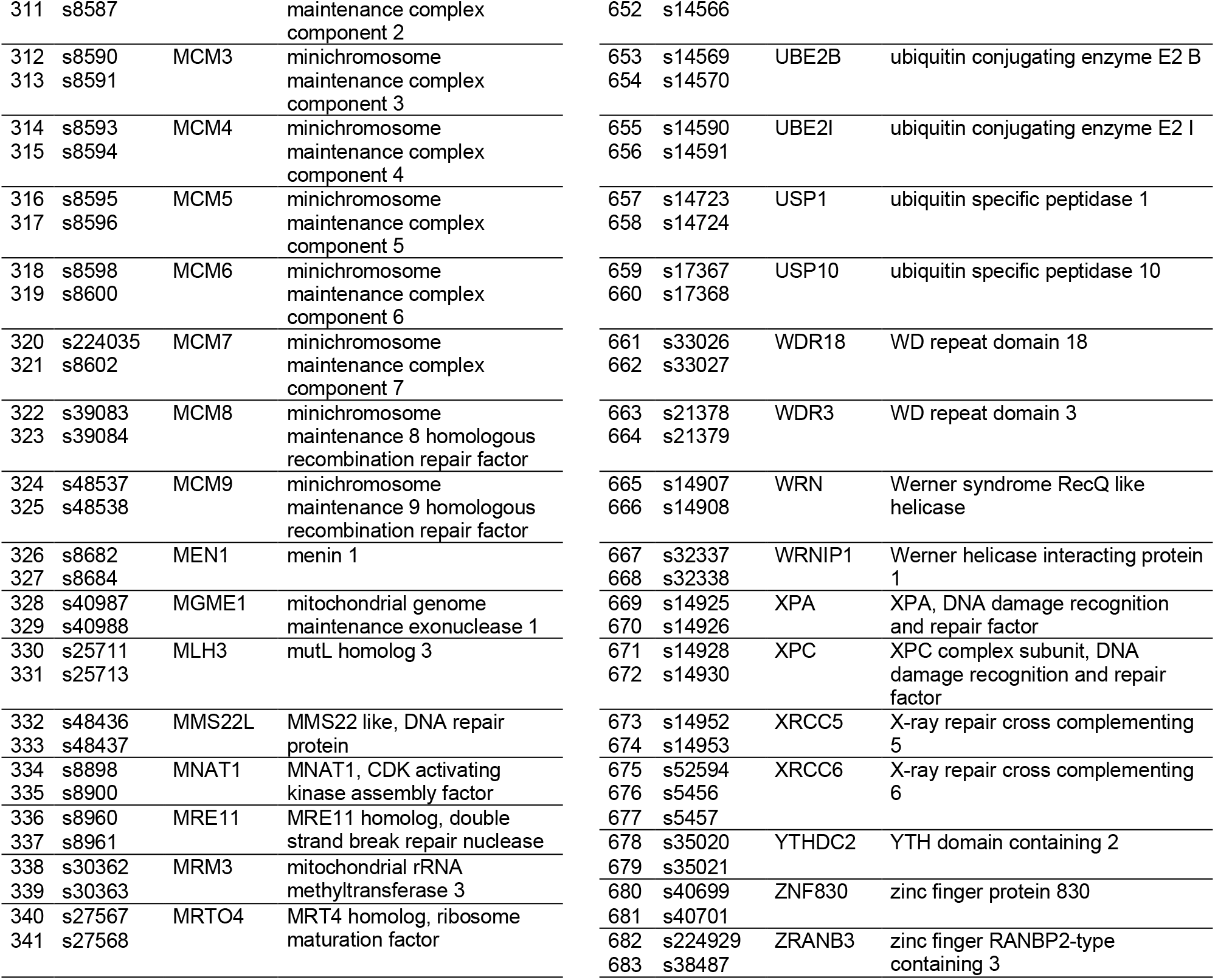
siRNA library of the primary screen.

**Supplementary Table 2.** Treatments/siRNAs included in the pilot screen.

| identifier | treatment type | target gene name | gene id | comment | control type |
| --- | --- | --- | --- | --- | --- |
| untreated | - | - | - | gelatin/sucrose coating only | negative |
| mock | transfection reagent | - | - | coating + transfection reagent | negative |
| sNEG1 | siRNA | - | - | non-targeting siRNA | negative |
| sNEG9 | siRNA | - | - | non-targeting siRNA | negative |
| sDEK | siRNA | DEK | DEK | - | transfection |
| APH | drug | - | - | 200 nM APH | positive |
| CPT | drug | - | - | 100 nM CPT | positive |
| sPOLD | siRNA | DNA polymerase $\delta$ | POLD1 | catalytic subunit | positive |
| sPOLA | siRNA | DNA polymerase $\alpha$ | POLA1 | catalytic subunit | positive |
| sKIF11 | siRNA | kinesin family member 11 | KIF11 | microtubule motor protein | phenotypic |
| sAURKB | siRNA | aurora kinase B | AURKB | binds to mitotic spindle | phenotypic |

**Supplementary Table 3.** Antibodies used in this study.

|  |  |  |
| --- | --- | --- |
| **Whole cell lysates:** |  |  |
| anti-DEK rabbit polyclonal K-877 | 1:20.000 | F. Kappes |
| anti-GFP mouse monoclonal | 1:1.000 | Roche (1 814 460) |
| Goat-anti-rabbit-HRP | 1:2.000 | Dako Denmark (P0448) |
| Goat-anti-mouse-HRP | 1:2.000 | Dako Denmark (P0447) |
| Streptavidin-biotinylated-HRP | 1:15.000 | GE Healthcare |
| **DEK-IP:** |  |  |
| anti-DEK mouse monoclonal | 1:500 | BD (610948) |
| anti-SUMO-2/3/4 mouse monoclonal Clone C-3 | 1:500 | Santa Cruz (sc-393144) |
| anti SUMO-1 mouse monoclonal Clone D-11 | 1:200 | Santa Cruz (sc-5308) |
| Veriblot for IP Detection Reagent (HRP) | 1:200 | Abcam (ab131366) |
| Streptavidin-biotinylated-HRP | 1:50.000 | GE Healthcare |
| **6His-SUMO Ni-NTA chromatography:** |  |  |
| anti-DEK rabbit polyclonal K-877 | 1:20.000 | F. Kappes |
| anti-SUMO-2/3/4 mouse monoclonal Clone C-3 | 1:500 | Santa Cruz (sc-393144) |
| anti SUMO-1 mouse monoclonal Clone D-11 | 1:200 | Santa Cruz (sc-5308) |
| Goat-anti-rabbit-HRP | 1:2.000 | Dako Denmark |
| Goat-anti-mouse-HRP | 1:2.000 | Dako Denmark |
| Streptavidin-biotinylated-HRP | 1:5.000 | GE Healthcare |
| **In vitro SUMOylation:** |  |  |
| anti-DEK rabbit polyclonal K-877 | 1:20.000 | F. Kappes |
| Anti-GST mouse monoclonal | 1:2.000 | Santa Cruz (sc-138) |
| anti-SUMO-2/3 rabbit monoclonal Clone 18H8 | 1:500 | Cell Signaling (#4971) |
| anti SUMO-1 mouse monoclonal Clone D-11 | 1:200 | Santa Cruz (sc-5308) |
| Goat-anti-rabbit-HRP | 1:2.000 | Dako Denmark |
| Goat-anti-mouse-HRP | 1:2.000 | Dako Denmark |
| Streptavidin-biotinylated-HRP | 1:25.000 | GE Healthcare |
| **Immunofluorescence MCF10A** |  |  |
| anti-DEK mouse | 1:50 | Santa Cruz (sc-136222) |
| anti-H3K9ac rabbit | 1:250 | Invitrogen (710293) |
| anti-H3K9me3 rabbit | 1:500 | Abcam (ab8898) |
| anti-H3K27me3 rabbit | 1:1000 | Abcam (ab192985) |
| anti-lamin A rabbit | 1:1000 | Abcam (ab253000) |
| anti-PCNA rabbit | 1:50 | Sigma-Aldrich (HPA030522) |
| anti-CENP-A rabbit | 1:400 | Cell Signaling (2186) |
| 2° anti-mouse Alexa Fluor 488 | 1:250 | Invitrogen (A11001) |
| 2° anti-rabbit Alexa Fluor 488 | 1:250 | Invitrogen (A11008) |
| 2° anti-rabbit Atto 532 | 1:200 | Rockland (611-152-122) |
| 2° anti-rabbit Atto 594 | 1:200 | Sigma-Aldrich (77671) |
| 2° FluoTag®-X2 IgG1 anti-Mouse Atto 647N | 1:500 | NanoTag Biotechnologies (N2002) |
| 2° anti-mouse IgG F(ab')<sub>2</sub>-goat Alexa Fluor 647 | 1:240 | ThermoFisher (A21237) |

**Supplementary Table 4.**
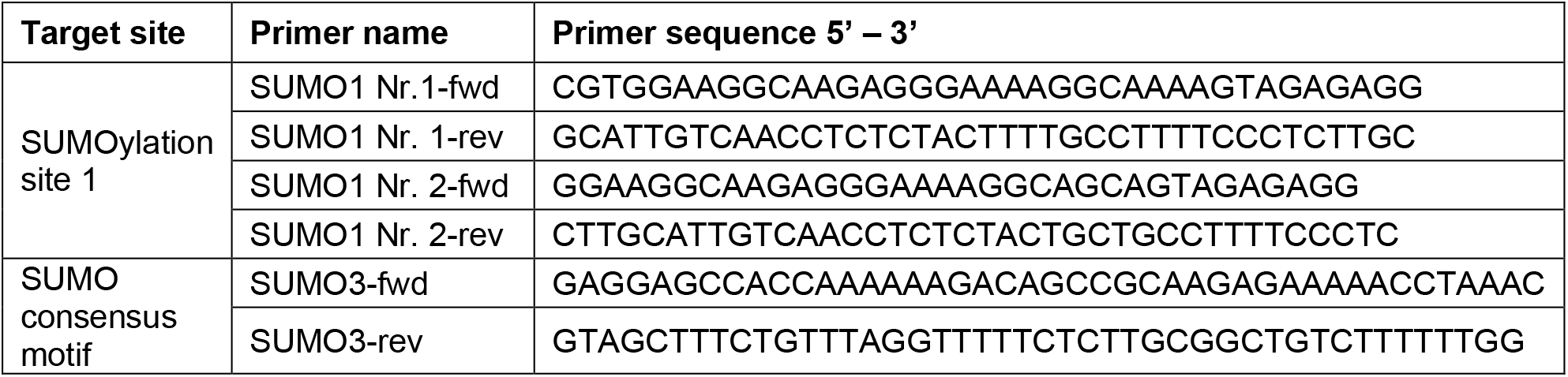
Primer sequences for the site-directed mutagenesis of the DEK primary sequence.

### Supplementary Materials and Methods

#### High-throughput siRNA screen coupled with time lapse fluorescence microscopy

U2-OS KI eGFP-DEK cells were transfected with siRNAs by solid-phase reverse transfection (Neumann et al., 2006): For the primary screen, the customized siRNA library comprised 678 unique siRNAs distributed on eight 96-well plates (the complete library can be found in **Supplementary Table 1**). In each plate, four wells contained the sNEG1 negative control, two wells the sPOLD1 (s616) positive control, two wells the sDEK transfection control (s15459) siRNA and three wells were left empty. The remaining wells contained the library siRNAs in alphabetical order. Before preparing the imaging plates, siRNAs of the mother plate were dissolved in 33 μl ddH_2_O to a final concentration of 3 μM and stored at -20 °C.

For the pilot screen, a 3 μM dilution of each control siRNA was distributed on a 96-well imaging plate (see **Supplementary Table 2**) in technical triplicates. Plates were sealed directly after distribution of siRNAs and stored at -20 °C until further use.

The following protocol was used for the coating of eight identical 96-well imaging plates from one unique siRNA layout. First, the siRNA plate was thawed and 5 μl of the 3 μM siRNA dilution were transferred into a V-shaped 96-well plate. Then, 11 μl of a mixture of 554 μl of 0.4 M sucrose in Opti-MEM, 323 μl of Lipofectamine 2000 and 323 μl of ddH2O were distributed into another 96-well plate. 7 μl of this transfection reagent mixture were transferred into the previously prepared siRNA plate and mixed 8 times. After 20 min of incubation, 7 μl of sterile filtered 0.2% gelatin (0.45 μm pore size) were added to the wells and mixed 8 times. This final transfection mix was diluted 1:51 in water in several steps. From this diluted transfection mix, 50 μl were distributed into each well of the 8 96-well COC imaging plates. In 100 μl medium the final siRNA concentration per well was 7.7 nM. Plates were immediately lyophilized (37 °C, 1 h) and stored in plastic boxes containing drying pearls.

#### Cell seeding and time-lapse microscopy

For the first four plates of the primary screen (layout no. 1-4, replicate 1), 4000 U2-OS GFP-DEK cells per well were seeded in 100 μl McCoy’s medium (without antibiotics) into the coated 96- well imaging plate using an automated cell seeding device and incubated for 48 h at 37 °C, 5% CO_2_. For all other plates, 3500 cells were seeded because cell confluency was slightly too high in some control wells. On the day of imaging, medium was exchanged with 100 μl pre warmed phenol red free CO_2_-independent imaging medium (Gibco) supplemented with 10% FBS, 1.5 mM L-glutamine and 2.3 g/l D-glucose. The full incubation IXM XLS high-content widefield microscope (Molecular Devices) was equipped with a PlanApochromat 20X/0.75 air objective and a sCMOS camera (4.66 megapixels). For GFP-DEK detection a 488 nm solid-state light source, the GFP filter set and a 690 nm laser autofocusing system were used while exposure time was set to 150 ms. For the time lapse experiment, four imaging field of views (FOVs) per well were set and images acquired every 12 min for 22 h. Two replicate plates per siRNA layout were seeded and imaged, for layout no. 5 three replicates. For the pilot screen, 2500 – 4000 cells per well were seeded, and 200 nM APH or 100 nM CPT were added to the imaging medium in the designated wells 1 h prior to imaging. For the time lapse experiment, 4-9 imaging FOVs per well were set and images acquired every 5 - 10 min for 24 - 48 h.

#### Image analysis using CellProfiler

The open-source software CellProfiler (v2.1.0 (Kamentsky et al., 2011)) was used for automated image analysis via a remote cluster to reduce computing time. The image analysis pipeline encompassed background correction, nuclei segmentation, filtering of dead and mitotic cells, intensity-based detection of DEK bodies and tracking of nuclei *via* the minimal overlap method.

The following steps were performed sequentially for every image of a time lapse. Images of the same time lapse were correlated using an object (here nuclei) tracking module. Then, the raw image was corrected for uneven illumination due to vignetting and for background fluorescence. As the effect was now limited to the corners of the image, the top and leftmost 30, the rightmost 60 and the bottommost 160 pixels (px) were cropped (“cropped image”). The cropped image was smoothed by applying a Gaussian filter and a threshold was calculated automatically with the global Otsu method to identify nuclei. Subsequently, the cropped image was masked with the nuclei outlines and the median intensity of the remaining background signal was subtracted from the cropped image (“corrected image”). Next, nuclei were segmented similarly by applying a Gaussian filter and calculating the threshold with the global Otsu method. Using this mask on the corrected image, nuclei of dead or mitotic cells were filtered out when meeting following requirements: Too high standard deviation and upper quartile of intensity and too small nuclear area. The filtered nuclei were tracked from image to the next image within a time lapse using a simple overlap method with a maximum pixel distance of 15. Afterwards, DEK bodies were identified: the corrected image was smoothed again by applying a Gaussian filter and DEK bodies were enhanced by performing a top-hat morphological operation. To avoid false-positive detection of peripheral micronuclei as DEK bodies, these protrusions were removed, and nuclei outlines were smoothed by performing an opening morphological operation. DEK bodies were identified within the smoothed outlines using a manual threshold (0.004 AU) and a diameter of 3 - 10 px. Finally, for each image CSV files containing image, nuclei and DEK body measurements were saved before the next image was analyzed. Aggregation of these raw data and evaluation of DEK body measurements was performed with the statistical software package R.

#### Data aggregation and transformation using R

The open-source programming language R with additional packages was used to aggregate the data. The goal was to obtain a single measurement per treatments/siRNA from the complex, time-resolved raw data of the technical (FOVs) and biological (plates) replicates that can be used to score DEK body regulators. As CellProfiler tracks the nuclei over subsequent frames/images, these track identifiers were used to summarize relevant measurements such as DEK body lifetime, number of DEK bodies per track etc. We noticed, however, that the CellProfiler pipeline not always managed to identify DEK bodies in all of the nuclei within a DEK body-containing sequence. We therefore wrote a custom function which used the closing morphological operation to fill these gaps with the DEK body counts of the previous or following frame (see lines 31-46 in the R code). In cases where a second, shorter DEK bodies-containing sequence was detected within the same track, the function also served to remove these likely false-negative DEK bodies by keeping only the longest sequence. Additionally, a similar function was implemented to calculate the mean DEK body number within this longest sequence, which was used as the main readout to score hits (see lines 50-60 in the R code). DEK body related measurements were then further aggregated over FOVs to investigate the variability within the same siRNAs/treatments. Afterwards, the aggregated data of all replicates was concatenated. For the primary screen, quality control was then performed based on the FOV summaries. FOVs with a small integrated nuclear fluorescence intensity (very small and/or dark nuclei), very low cell number, blurry images and a negative slope of their linear growth function were filtered out. Furthermore, all wells that were contaminated with bacteria or yeast after the time lapse (as seen under a brightfield benchtop microscope) were filtered out manually.

Lastly, Z-scores of the FOV-based summaries of the mean number of DEK bodies were calculated. The Z-score can be used when the data distribution is unimodal and variability of the control measurement (here sNEG1) does not change in between replicates (Zhang, 2011). Therefore, the data was log (base 2)-transformed and the mean (X_sNEG1_) and the standard deviation (SD_sNEG1_) of all negative control (sNEG1) FOVs from the same replicate were calculated. For all FOVs of the other siRNAs of this replicate a Z-score was computed:

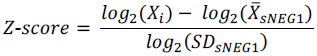

The mean of all Z-scores of the same siRNA was finally calculated. By ordering this mean DEK body number by size it was possible to score DEK body up-regulators (highest Z-score) and down-regulators (lowest Z-score). R code provided at https://github.com/CP-Vogel/R-DEK-body-script.git.

#### Low-throughput siRNA validation screen coupled with time-lapse fluorescence microscopy

For siRNA delivery, a liquid-phase instead of a solid-phase reverse transfection procedure was employed. Transfection mixes were freshly prepared for each replicate before cell seeding. Validation siRNAs were picked from the primary screen mother plates into a V-shaped 96-well plate and diluted to 100 nM with MQ H_2_O. Two wells were supplied with the same validation siRNA, four wells with sNEG1 negative control, two wells with sDEK transfection control (s15459) siRNA and four wells were left empty. The new mother plate was stored at -20 °C until the day of transfection. At the day of transfection, 8 μl of a 1:20 dilution of Lipofectamine RNAiMAX in Opti-MEM was added in each well of a 96-well imaging plate (Ibidi). 25 μl of diluted siRNAs were mixed with the pre-distributed transfection reagent dilution. After 15 min incubation, 5500 U2-OS GFP- DEK cells in 207 μl medium (no antibiotics) were added for a final siRNA concentration of 10.4 nM and cells were incubated for 48 h at 37°C, 5% CO2.

48 h after transfection, medium was exchanged with 200 μl of pre-warmed phenol red free imaging medium (Hyclone, no antibiotics) and equilibrated in a pre-warmed CellObserver HSl microscope (Zeiss) equipped with a CSU-X1 spinning disc unit (Yokogawa) and a full incubator for 1 h at 37 °C, 5% CO2, humid atmosphere. Images were acquired with a PlanApochromat 20X/0.8 air objective, a 488 nm OPSL laser for GFP-DEK detection and an Evolve EMCCD camera (Photometrics, 512 x 512 pixels). Exposure time and EM gain were set to 150 ms and 700, respectively. For the time-lapse experiment, 6 imaging FOVs per well were set, the focus adjusted manually and images acquired every 12 min for 24 h. Three replicate plates were acquired in total.

In contrast to the primary screen, the image analysis was run on a newer version of CellProfiler (v.2.2.0 instead of 2.1.0) and locally on a workstation instead of a remote cluster. The pipeline was adjusted from the primary screen to meet the needs of the new microscope setup. Raw data analysis was performed as for the primary screen with minor modifications. Calculation of Z-scores and siRNA summaries was performed as described above.

## References

1. Abbas, T. (2021). The Role of Ubiquitination and SUMOylation in DNA Replication. Curr. Issues Mol. Biol. 40, 189–220.

2. Aichem, A., Sailer, C., Ryu, S., Catone, N., Stankovic-Valentin, N., Schmidtke, G., Melchior, F., Stengel, F. and Groettrup, M. (2019). The ubiquitin-like modifier FAT10 interferes with SUMO activation. Nat. Commun. 10,.

3. Alabert, C., Bukowski-Wills, J. C., Lee, S. B., Kustatscher, G., Nakamura, K., De Lima Alves, F., Menard, P., Mejlvang, J., Rappsilber, J. and Groth, A. (2014). Nascent chromatin capture proteomics determines chromatin dynamics during DNA replication and identifies unknown fork components. Nat. Cell Biol. 16, 281–291.

4. Aladjem, M. I. (2007). Replication in context: Dynamic regulation of DNA replication patterns in metazoans. Nat. Rev. Genet. 8, 588–600.

5. Alexiadis, V., Waldmann, T., Andersen, J., Mann, M., Knippers, R. and Gruss, C. (2000). The protein encoded by the proto-oncogene DEK changes the topology of chromatin and reduces the efficiency of DNA replication in a chromatin specific manner. Genes Dev. 14, 1308–1312.

6. Aranda, S., Rutishauser, D. and Ernfors, P. (2014). Identification of a large protein network involved in epigenetic transmission in replicating DNA of embryonic stem cells. Nucleic Acids Res. 42, 6972–6986.

7. Baltz, A. G., Munschauer, M., Schwanhäusser, B., Vasile, A., Murakawa, Y., Schueler, M., Youngs, N., Penfold-Brown, D., Drew, K., Milek, M., et al. (2012). The mRNA-bound proteome and its global occupancy profile on protein-coding transcripts. Mol. Cell 46, 674–690.

8. Bates, M., Huang, B., Dempsey, G. T. and Zhuang, X. (2007). Multicolor super-resolution imaging with photo-switchable fluorescent probes. Science 317, 1749–53.

9. Beckmann, B. M., Horos, R., Fischer, B., Castello, A., Eichelbaum, K., Alleaume, A.-M., Schwarzl, T., Curk, T., Foehr, S., Huber, W., et al. (2015). The RNA-binding proteomes from yeast to man harbour conserved enigmRBPs. Nat. Commun. 6, 10127.

10. Bergoglio, V., Boyer, A. S., Walsh, E., Naim, V., Legube, G., Lee, M. Y. W. T., Rey, L., Rosselli, F., Cazaux, C., Eckert, K. A., et al. (2013). DNA synthesis by pol η promotes fragile site stability by preventing under-replicated DNA in mitosis. J. Cell Biol. 201, 395–408.

11. Bhat, A., Andersen, P. L., Qin, Z. and Xiao, W. (2013). Rev3, the catalytic subunit of Polζ, is required for maintaining fragile site stability in human cells. Nucleic Acids Res. 41, 2328–2339.

12. Böhm, F., Kappes, F., Scholten, I., Richter, N., Matsuo, H., Knippers, R. and Waldmann, T. (2005). The SAF-box domain of chromatin protein DEK. Nucleic Acids Res. 33, 1101–1110.

13. Boyer, A. S., Walter, D. and Sørensen, C. S. (2016). DNA replication and cancer: From dysfunctional replication origin activities to therapeutic opportunities. Semin. Cancer Biol. 37–38, 16–25.

14. Capitano, M. L., Markovitz, D. M., Hal, E., Invest, J. C., Capitano, M. L., Mor-vaknin, N., Saha, A. K., Cooper, S., Legendre, M., Guo, H., et al. (2019). Secreted nuclear protein DEK regulates hematopoiesis through CXCR2 signaling Find the latest version : Secreted nuclear protein DEK regulates hematopoiesis through CXCR2 signaling. 129, 2555–2570.

15. Capitano, M. L., Sammour, Y., Ropa, J., Legendre, M., Mor-Vaknin, N. and Markovitz, D. M. (2022). DEK, a nuclear protein, is chemotactic for hematopoietic stem/progenitor cells acting through CXCR2 and Gαi signaling. J. Leukoc. Biol. 112, 449–456.

16. Casas-Delucchi, C. S., Van Bemmel, J. G., Haase, S., Herce, H. D., Nowak, D., Meilinger, D., Stear, J. H., Leonhardt, H. and Cardoso, M. C. (2012). Histone hypoacetylation is required to maintain late replication timing of constitutive heterochromatin. Nucleic Acids Res. 40, 159–169.

17. Castello, A., Fischer, B., Eichelbaum, K., Horos, R., Beckmann, B. M., Strein, C., Davey, N. E., Humphreys, D. T., Preiss, T., Steinmetz, L. M., et al. (2012). Insights into RNA biology from an atlas of mammalian mRNA-binding proteins. Cell 149, 1393–1406.

18. Cavellán, E., Asp, P., Percipalle, P. and Farrants, A. K. Ö. (2006). The WSTF-SNF2h chromatin remodeling complex interacts with several nuclear proteins in transcription. J. Biol. Chem. 281, 16264–16271.

19. Cerase, A., Pintacuda, G., Tattermusch, A. and Avner, P. (2015). Xist localization and function: New insights from multiple levels. Genome Biol. 16, 1–12.

20. Chadwick, B. P. and Willard, H. F. (2004). Multiple spatially distinct types of facultative heterochromatin on the human inactive X chromosome. Proc. Natl. Acad. Sci. U. S. A. 101, 17450–17455.

21. Chagin, V. O., Casas-Delucchi, C. S., Reinhart, M., Schermelleh, L., Markaki, Y., Maiser, a, Bolius, J. J., Bensimon, a, Fillies, M., Domaing, P., et al. (2016). 4D Visualization of replication foci in mammalian cells corresponding to individual replicons. Nat. Commun. 7, 11231.

22. Cleary, J., Sitwala, K. V., Khodadoust, M. S., Kwok, R. P. S., Mor-Vaknin, N., Cebrat, M., Cole, P. A. and Markovitz, D. M. (2005). p300/CBP-associated factor drives DEK into interchromatin granule clusters. J. Biol. Chem. 280, 31760–31767.

23. Despras, E., Sittewelle, M., Pouvelle, C., Delrieu, N., Cordonnier, A. M. and Kannouche, P. L. (2016). Rad18-dependent SUMOylation of human specialized DNA polymerase eta is required to prevent under-replicated DNA. Nat. Commun. 7, 13326.

24. Deutzmann, A., Ganz, M., Schönenberger, F., Vervoorts, J., Kappes, F. and Ferrando-May,E. (2015). The human oncoprotein and chromatin architectural factor DEK counteracts DNA replication stress. Oncogene 34, 4270–4277.

25. Diaspro, A. and Bianchini, P. (2020). Optical nanoscopy. Riv. del Nuovo Cim. 43, 385–455.

26. Dundr, M. (2012). Nuclear bodies: multifunctional companions of the genome. Curr. Opin. Cell Biol. 24, 415–422.

27. Fahrer, J., Popp, O., Malanga, M., Beneke, S., Markovitz, D. M., Ferrando-May, E., Bürkle, A. and Kappes, F. (2010). High-affinity interaction of poly(ADP-ribose) and the human DEK oncoprotein depends upon chain length. Biochemistry 49, 7119–7130.

28. Fragkos, M., Ganier, O., Coulombe, P. and Méchali, M. (2015). DNA replication origin activation in space and time. Nat. Rev. Mol. Cell Biol. 16, 360–374.

29. Gamble, M. J. and Fisher, R. P. (2007). SET and PARP1 remove DEK from chromatin to permit access by the transcription machinery. Nat. Struct. Mol. Biol. 14, 548–555.

30. Ganz, M., Vogel, C., Czada, C., Jörke, V., Gwosch, E. C., Kleiner, R., Pierzynska-Mach, A., Cella Zanacchi, F., Diaspro, A., Kappes, F., et al. (2019). The oncoprotein DEK affects the outcome of PARP1/2 inhibition during mild replication stress. PLoS One 14, e0213130.

31. Garcia, P. A. A., Hoover, M. E., Zhang, P., Nagarajan, P., Freitas, M. A. and Parthun, M. R. (2017). Identification of multiple roles for histone acetyltransferase 1 in replication-coupled chromatin assembly. Nucleic Acids Res. 45, 9319–9335.

32. Grant, M. M. (2010). Identification of SUMOylated proteins in neuroblastoma cells after treatment with hydrogen peroxide or ascorbate. BMB Rep. 43, 720–725.

33. Guérillon, C., Smedegaard, S., Hendriks, I. A., Nielsen, M. L. and Mailand, N. (2020). Multisite SUMOylation restrains DNA polymerase η interactions with DNA damage sites. J. Biol. Chem. 295, 8350–8362.

34. Guo, H., Prell, M., Königs, H., Xu, N., Waldmann, T., Hermans-Sachweh, B., Ferrando-May, E., Lüscher, B. and Kappes, F. (2021a). Bacterial Growth Inhibition Screen (BGIS) identifies a loss-of-function mutant of the DEK oncogene, indicating DNA modulating activities of DEK in chromatin. FEBS Lett. 595, 1438–1453.

35. Guo, H., Xu, N., Prell, M., Königs, H., Hermanns-Sachweh, B., Lüscher, B. and Kappes, F. (2021b). Bacterial Growth Inhibition Screen (BGIS): harnessing recombinant protein toxicity for rapid and unbiased interrogation of protein function. FEBS Lett. 595, 1422–1437.

36. Heinz, K. S., Casas-Delucchi, C. S., Török, T., Cmarko, D., Rapp, A., Raska, I. and Cardoso, M. C. (2018). Peripheral re-localization of constitutive heterochromatin advances its replication timing and impairs maintenance of silencing marks. Nucleic Acids Res. 46, 6112–6128.

37. Hollenbach, A. D., McPherson, C. J., Mientjes, E. J., Iyengar, R. and Grosveld, G. (2002). Daxx and histone deacetylase II associate with chromatin through an interaction with core histones and the chromatin-associated protein Dek. J. Cell Sci. 115, 3319–3330.

38. Hu, H. G., Illges, H., Gruss, C. and Knippers, R. (2005). Distribution of the chromatin protein DEK distinguishes active and inactive CD21/CR2 gene in pre and mature B lymphocytes. Int. Immunol. 17, 789–796.

39. Hu, H. G., Scholten, I., Gruss, C. and Knippers, R. (2007). The distribution of the DEK protein in mammalian chromatin. Biochem. Biophys. Res. Commun. 358, 1008–1014.

40. Ivanauskiene, K., Delbarre, E., McGhie, J. D., Kuntziger, T., Wong, L. H. and Collas, P. (2014). The PML-associated protein DEK regulates the balance of H3.3 loading on chromatin and is important for telomere integrity. Genome Res. 24, 1584–1594.

41. Jackson, D. A. and Pombo, A. (1998). Replicon clusters are stable units of chromosome structure: Evidence that nuclear organization contributes to the efficient activation and propagation of S phase in human cells. J. Cell Biol. 140, 1285–1295.

42. Kamentsky, L., Jones, T. R., Fraser, A., Bray, M. A., Logan, D. J., Madden, K. L., Ljosa, V., Rueden, C., Eliceiri, K. W. and Carpenter, A. E. (2011). Improved structure, function and compatibility for cellprofiler: Modular high-throughput image analysis software. Bioinformatics 27, 1179–1180.

43. Kappes, F., Burger, K., Baack, M., Fackelmayer, F. O. and Gruss, C. (2001). Subcellular Localization of the Human Proto-oncogene Protein DEK. J. Biol. Chem. 276, 26317–26323.

44. Kappes, F., Damoc, C., Knippers, R., Przybylski, M., Pinna, L. A. and Gruss, C. (2004a). Phosphorylation by Protein Kinase CK2 Changes the DNA Binding Properties of the Human Chromatin Protein DEK. Mol. Cell. Biol. 24, 6011–6020.

45. Kappes, F., Scholten, I., Richter, N., Gruss, C. and Waldmann, T. (2004b). Functional Domains of the Ubiquitous Chromatin Protein DEK. Mol. Cell. Biol. 24, 6000–6010.

46. Kappes, F., Fahrer, J., Khodadoust, M. S., Tabbert, A., Strasser, C., Mor-Vaknin, N., Moreno-Villanueva, M., Burkle, A., Markovitz, D. M. and Ferrando-May, E. (2008). DEK Is a Poly(ADP-Ribose) Acceptor in Apoptosis and Mediates Resistance to Genotoxic Stress. Mol. Cell. Biol. 28, 3245–3257.

47. Kappes, F., Waldmann, T., Mathew, V., Yu, J., Zhang, L., Khodadoust, M. S., Chinnaiyan, A. M., Luger, K., Erhardt, S., Schneider, R., et al. (2011a). The DEK oncoprotein is a Su(var) that is essential to heterochromatin integrity. Genes Dev. 25, 673–678.

48. Kappes, F., Khodadoust, M. S., Yu, L., Kim, D. S. L., Fullen, D. R., Markovitz, D. M. and Ma, L. (2011b). DEK expression in melanocytic lesions. Hum. Pathol. 42, 932–938.

49. Karam, M., Thenoz, M., Capraro, V., Robin, J. P., Pinatel, C., Lançon, A., Galia, P., Sibon, D., Thomas, X., Ducastelle-Lepretre, S., et al. (2014). Chromatin redistribution of the DEK oncoprotein represses hTERT transcription in leukemias. Neoplasia (United States) 16, 21–30.

50. Lossaint, G., Larroque, M., Ribeyre, C., Bec, N., Larroque, C., Décaillet, C., Gari, K. and Constantinou, A. (2013). FANCD2 Binds MCM Proteins and Controls Replisome Function upon Activation of S Phase Checkpoint Signaling. Mol. Cell 51, 678–690.

51. Lukusa, T. and Fryns, J. P. (2008). Human chromosome fragility. Biochim. Biophys. Acta - Gene Regul. Mech. 1779, 3–16.

52. Mailand, N., Gibbs-Seymour, I. and Bekker-Jensen, S. (2013). Regulation of PCNA-protein interactions for genome stability. Nat. Rev. Mol. Cell Biol. 14, 269–282.

53. Matrka, M. C., Hennigan, R. F., Kappes, F., Delay, M. L., Lambert, P. F., Aronow, B. J. and Wells, S. I. (2015). DEK over-expression promotes mitotic defects and micronucleus formation. Cell Cycle 14, 3939–3953.

54. Maya-Mendoza, A., Moudry, P., Merchut-Maya, J. M., Lee, M., Strauss, R. and Bartek, J. (2018). High speed of fork progression induces DNA replication stress and genomic instability. Nature 559, 279–284.

55. Mintz, P. J., Patterson, S. D., Neuwald, A. F., Spahr, C. S. and Spector, D. L. (1999). Purification and biochemical characterization of interchromatin granule clusters. EMBO J. 18, 4308–4320.

56. Mirkin, E. V. and Mirkin, S. M. (2007). Replication Fork Stalling at Natural Impediments. Microbiol. Mol. Biol. Rev. 71, 13–35.

57. Mor-Vaknin, N., Kappes, F., Dick, A. E., Legendre, M., Damoc, C., Teitz-Tennenbaum, S., Kwok, R., Ferrando-May, E., Adams, B. S. and Markovitz, D. M. (2011). DEK in the synovium of patients with juvenile idiopathic arthritis: characterization of DEK antibodies and posttranslational modification of the DEK autoantigen. Arthritis Rheum. 63, 556–567.

58. Nakayama, J., Rice, J. C., Strahl, B. D., Allis, C. D. and Grewal, S. I. S. (2001). Role of Histone H3 Lysine 9 Methylation in Epigenetic Control of Heterochromatin Assembly. Science (80-). 292, 110–113.

59. Neumann, B., Held, M., Liebel, U., Erfle, H., Rogers, P., Pepperkok, R. and Ellenberg, J. (2006). High-throughput RNAi screening by time-lapse imaging of live human cells. Nat. Methods 3, 385–390.

60. O’Keefe, R. T., Henderson, S. C. and Spector, D. L. (1992). Dynamic organization of DNA replication in mammalian cell nuclei: Spatially and temporally defined replication of chromosome-specific α- satellite DNA sequences. J. Cell Biol. 116, 1095–1110.

61. Özçelik, E., Kalaycı, A., Çelik, B., Avcı, A., Akyol, H., Kılıç, İ. B., Güzel, T., Çetin, M., Öztürk, M. T., Çalışkaner, Z. O., et al. (2022). Doxorubicin induces prolonged DNA damage signal in cells overexpressing DEK isoform-2. PLoS One 17, e0275476.

62. Penagos-Puig, A. and Furlan-Magaril, M. (2020). Heterochromatin as an Important Driver of Genome Organization. Front. Cell Dev. Biol. 8, 1–10.

63. Privette Vinnedge, L. M., Kappes, F., Nassar, N. and Wells, S.I. (2013). Stacking the DEK: From chromatin topology to cancer stem cells. Cell Cycle 12, 51–66.

64. Ribeyre, C., Zellweger, R., Chauvin, M., Bec, N., Larroque, C., Lopes, M. and Constantinou, A. (2016). Nascent DNA Proteomics Reveals a Chromatin Remodeler Required for Topoisomerase I Loading at Replication Forks. Cell Rep. 15, 300–309.

65. Ryba, T., Hiratani, I., Lu, J., Itoh, M., Kulik, M., Zhang, J., Schulz, T. C., Robins, A. J., Dalton, S. and Gilbert, D. M. (2010). Evolutionarily conserved replication timing profiles predict long range chromatin interactions and distinguish closely related cell types. Genome Res. 20, 761–770.

66. Saha, A. K., Kappes, F., Mundade, A., Deutzmann, A., Rosmarin, D. M., Legendre, M., Chatain, N., Al-Obaidi, Z., Adams, B. S., Ploegh, H. L., et al. (2013). Intercellular trafficking of the nuclear oncoprotein DEK. Proc. Natl. Acad. Sci. U. S. A. 110, 6847–52.

67. Sandén, C. and Gullberg, U. (2015). The DEK oncoprotein and its emerging roles in gene regulation. Leukemia 29, 1632–1636.

68. Sandén, C., Järvstråt, L., Lennartsson, A., Brattås, P. L., Nilsson, B. and Gullberg, U. (2014). The DEK oncoprotein binds to highly and ubiquitously expressed genes with a dual role in their transcriptional regulation. Mol. Cancer 13, 1–13.

69. Sawatsubashi, S., Murata, T., Lim, J., Fujiki, R., Ito, S., Suzuki, E., Tanabe, M., Zhao, Y., Kimura, S., Fujiyama, S., et al. (2010). A histone chaperone, DEK, transcriptionally coactivates a nuclear receptor. Genes Dev. 24, 159–170.

70. Saxena, S. and Zou, L. (2022). Hallmarks of DNA replication stress. Mol. Cell 82, 2298–2314.

71. Sehwaiger, M., Stadler, M. B., Bell, O., Kohler, H., Oakeley, E. J. and Sehübeler, D. (2009). Chromatin state marks cell-type and gender-specific replication of the Drosophila genome. Genes Dev. 23, 589–601.

72. Sirbu, B. M., McDonald, W. H., Dungrawala, H., Badu-Nkansah, A., Kavanaugh, G. M., Chen, Y., Tabb, D. L. and Cortez, D. (2013). Identification of proteins at active, stalled, and collapsed replication forks using isolation of proteins on nascent DNA (iPOND) coupled with mass spectrometry. J. Biol. Chem. 288, 31458–31467.

73. Smith, E. A., Krumpelbeck, E. F., Jegga, A. G., Prell, M., Matrka, M. M., Kappes, F., Greis, K. D., Ali, A. M., Meetei, A. R. and Wells, S. I. (2018). The nuclear DEK interactome supports multi-functionality “The DEK Interactome.” Proteins Struct. Funct. Bioinforma. 86, 88–97.

74. Stepanenko, O. V, Sulatsky, M. I., Mikhailova, E. V, Kuznetsova, I. M., Turoverov, K. K., Stepanenko, O. V and Sulatskaya, A. I. (2021). New findings on GFP-like protein application as fluorescent tags: Fibrillogenesis, oligomerization, and amorphous aggregation. Int. J. Biol. Macromol. 192, 1304–1310.

75. Stewart-Morgan, K. R., Petryk, N. and Groth, A. (2020). Chromatin replication and epigenetic cell memory. Nat. Cell Biol. 22, 361–371.

76. Su, Q. P., Zhao, Z. W., Meng, L., Ding, M., Zhang, W., Li, Y., Liu, M., Li, R., Gao, Y. Q., Xie, X. S., et al. (2020). Superresolution imaging reveals spatiotemporal propagation of human replication foci mediated by CTCF-organized chromatin structures. Proc. Natl. Acad. Sci. U.S. A. 117, 15036–15046.

77. Tabbert, A., Kappes, F., Knippers, R., Kellermann, J., Lottspeich, F. and Ferrando-May, E. (2006). Hypophosphorylation of the architectural chromatin protein DEK in death-receptor induced apoptosis revealed by the isotope coded protein label proteomic platform. Proteomics 6, 5758–5772.

78. Tatham, M. H., Rodriguez, M. S., Xirodimas, D. P. and Hay, R. T. (2009). Detection of protein SUMOylation in vivo. Nat. Protoc. 4, 1363–1371.

79. Trojer, P. and Reinberg, D. (2007). Facultative Heterochromatin: Is There a Distinctive Molecular Signature? Mol. Cell 28, 1–13.

80. Tsao, W.-C. and Eckert, K. A. (2018). Detours to Replication: Functions of Specialized DNA Polymerases during Oncogene-induced Replication Stress. Int. J. Mol. Sci. 19,.

81. Vesela, E., Chroma, K., Turi, Z. and Mistrik, M. (2017). Common Chemical Inductors of Replication Stress: Focus on Cell-Based Studies. Biomolecules 7, 19.

82. Waidmann, S., Kusenda, B., Mayerhofer, J., Mechtler, K. and Jonak, C. (2014). A DEK domain-containing protein modulates chromatin structure and function in Arabidopsis. Plant Cell 26, 4328–4344.

83. Waldmann, T., Eckerich, C., Baack, M. and Gruss, C. (2002). The ubiquitous chromatin protein DEK alters the structure of DNA by introducing positive supercoils. J. Biol. Chem. 277, 24988–24994.

84. Waldmann, T., Baack, M., Richter, N. and Gruss, C. (2003). Structure-specific binding of the proto-oncogene protein DEK to DNA. Nucleic Acids Res. 31, 7003–7010.

85. Waldmann, T., Scholten, I., Kappes, F., Hu, H. G. and Knippers, R. (2004). The DEK protein an abundant and ubiquitous constituent of mammalian chromatin. Gene 343, 1–9.

86. Wang, Z., Zang, C., Rosenfeld, J. A., Schones, D. E., Barski, A., Cuddapah, S., Cui, K., Roh, T. Y., Peng, W., Zhang, M. Q., et al. (2008). Combinatorial patterns of histone acetylations and methylations in the human genome. Nat. Genet. 40, 897–903.

87. Wessel, D. and Flügge, U. I. (1984). A method for the quantitative recovery of protein in dilute solution in the presence of detergents and lipids. Anal. Biochem. 138, 141–143.

88. Yang, Y.-S., Jia, X.-Z., Lu, Q.-Y., Cai, S.-L., Huang, X.-T., Yang, S.-H., Wood, C., Wang, Y.-H., Zhou, J.-J., Chen, Y.-D., et al. (2022). Exosomal DEK removes chemoradiotherapy resistance by triggering quiescence exit of breast cancer stem cells. Oncogene 41, 2624–2637.

89. Zhang, X. D. (2011). Illustration of SSMD, z score, SSMD*, z* score, and t statistic for hit selection in RNAi high-throughput screens. J. Biomol. Screen. 16, 775–785.

90. Zhang, Q., Bassetti, F., Gherardi, M. and Lagomarsino, M. C. (2017). Cell-to-cell variability and robustness in S-phase duration from genome replication kinetics. Nucleic Acids Res. 45, 8190–8198.

91. Zhao, Q., Xie, Y., Zheng, Y., Jiang, S., Liu, W., Mu, W., Liu, Z., Zhao, Y., Xue, Y. and Ren, J. (2014). GPS-SUMO: A tool for the prediction of sumoylation sites and SUMO-interaction motifs. Nucleic Acids Res. 42, 325–330.

92. Zheng, L., Baumann, U. and Reymond, J. L. (2004). An efficient one-step site-directed and site saturation mutagenesis protocol. Nucleic Acids Res. 32,.

93. Zhou, J., Zhao, L., Wu, Y., Zhang, X., Cheng, S., Wei, F., Zhang, Y., Zhu, H., Zhou, Y., Feng, Z., et al. (2022). A DEK domain-containing protein GhDEK2D mediated Gossypium hirsutum enhanced resistance to Verticillium dahliae. Plant Signal. Behav. 17,

